# PETRA: Prime editing of transcribed regulatory elements to assay expression

**DOI:** 10.1101/2025.11.18.689114

**Authors:** Magdalena Armas Reyes, Michael Herger, Laura Cubitt, Gregory M. Findlay

**Author notes:** Correspondence to Gregory M. Findlay.

## Abstract

Predicting how changes in human DNA sequence impact gene expression remains challenging. Here, we present PETRA (<u>P</u>rime <u>E</u>diting of <u>T</u>ranscribed <u>R</u>egulatory elements to <u>A</u>ssay expression), a multiplexed genome editing method to quantify the effects of regulatory variants at scale. PETRA leverages the delivery of variants to abundantly transcribed regions of genes such that sequence-specific effects on mRNA expression can be read out by amplicon sequencing. We demonstrate PETRA in Jurkat cells by scoring 13,935 six-nucleotide insertions delivered to the 5’ untranslated regions (5’ UTRs) of four genes important for T cell responses, namely *VAV1*, *IL2RA*, *CD28* and *OTUD7B*. Effects on expression are linked to the creation of new transcription factor binding sites (TFBSs), as well as to alterations in splicing and translation initiation. Combinatorial delivery of TFBSs identified using PETRA generates alleles that increase mRNA expression more than 10-fold. Additionally, we extend PETRA to primary human T cells to compare effects across cell types and use our data to assess the performance of computational models. These results establish PETRA as a flexible means of dissecting and reprogramming the logic of gene regulation across genomic contexts and cell types.

## INTRODUCTION

Identification of DNA sequence variants impacting human gene expression drives biological discovery and improves the interpretation of personal genomes. Beyond understanding which variants impact human phenotypes, knowledge of gene regulation can also be directly leveraged for therapeutic benefit. For example, genome editing of regulatory elements has emerged as a means of treating genetic disease^1^. While altering gene expression by editing the genome has a broad range of potential applications, such strategies are currently limited by our inability to accurately predict how specific changes in DNA sequence lead to changes in gene expression.

Computational models that predict expression from DNA sequence have improved substantially in recent years, but still perform poorly at predicting effects of specific variants^2–8^. One reason for this is that such models are hindered by a lack of high-quality training data. Large-scale omics studies have succeeded at associating common variants with changes in gene expression, chromatin states, and phenotypes^9–11^. However, such observational approaches are of limited benefit for inferring causal effects of new variants. Meanwhile, high-throughput experimental systems, such as massively parallel reporter assays (MPRAs), have enabled the interrogation of regulatory elements and variants for expression effects at scale^12,13^. Though valuable, MPRAs make use of artificial reporter cassettes that are expressed exogenously or via random genomic integration^14^. Thus, key features of variants’ native loci are lost, such as surrounding DNA sequence and chromatin context. To date, relatively few variants have been engineered in their native context to assay expression effects, either in immortalised lines or primary cells.

While CRISPR-based methods have been established to assay effects of variants, all have limitations. CRISPR interference (CRISPRi) and CRISPR activation (CRISPRa) technologies enable identification of noncoding elements of functional importance^15–18^, but do not probe the effects of sequence variants. CRISPR and base editing screens^15,19^ afford the ability to assay large libraries but are restricted to limited types of variants (i.e., indels or specific substitutions). Meanwhile, saturation genome editing (SGE) has proven clinically valuable for assaying coding variants in cell lines suited to editing by homology-directed DNA repair^20^, but has yet to show broad utility for regulatory variants. Very recently, multiplex prime editing (PE) strategies have been reported to assay variants for effects on expression^21,22^. However, thus far such methods have only been applied to immortalised cell lines and scalability has been limited by the need to develop a FACS-based read-out for each target gene. More broadly applicable, high-throughput methods to experimentally test regulatory variants in their native context will prove highly valuable.

Here, we introduce <u>P</u>rime <u>E</u>diting of <u>T</u>ranscribed <u>R</u>egions to <u>A</u>ssay expression, or PETRA. We demonstrate PETRA by assaying a total of 14,888 variants across four genes with roles in human T cell responses, successfully identifying individual edits, combinations of edits, and transcription factor binding sites (TFBSs) that increase messenger RNA (mRNA) expression by greater than 10-fold in Jurkat and primary human T cells. We anticipate application of PETRA to diverse loci and cell types will refine our understanding of gene expression, facilitate rapid accumulation of data for benchmarking and training computational models, and speed the discovery of edits that confer therapeutic benefit.

## RESULTS

### Prime Editing of Transcribed Regulatory Elements to Assay expression

With the aim of measuring endogenously engineered variants’ effects on gene expression in high-throughput, we developed PETRA. PETRA leverages PE^23^ to install variants in non-coding regions of target genes and next-generation sequencing (NGS) to readout their effects on expression (**Figure 1A**). We reasoned that by editing abundantly transcribed regions, regulatory effects of variants could be determined in multiplex simply by performing amplicon sequencing of RNA and DNA from edited cells. In turn, this would allow individual variants and combinations of variants to be read out in an allele-specific manner, potentially across each of many target sites and cell types.

**Figure 1.**
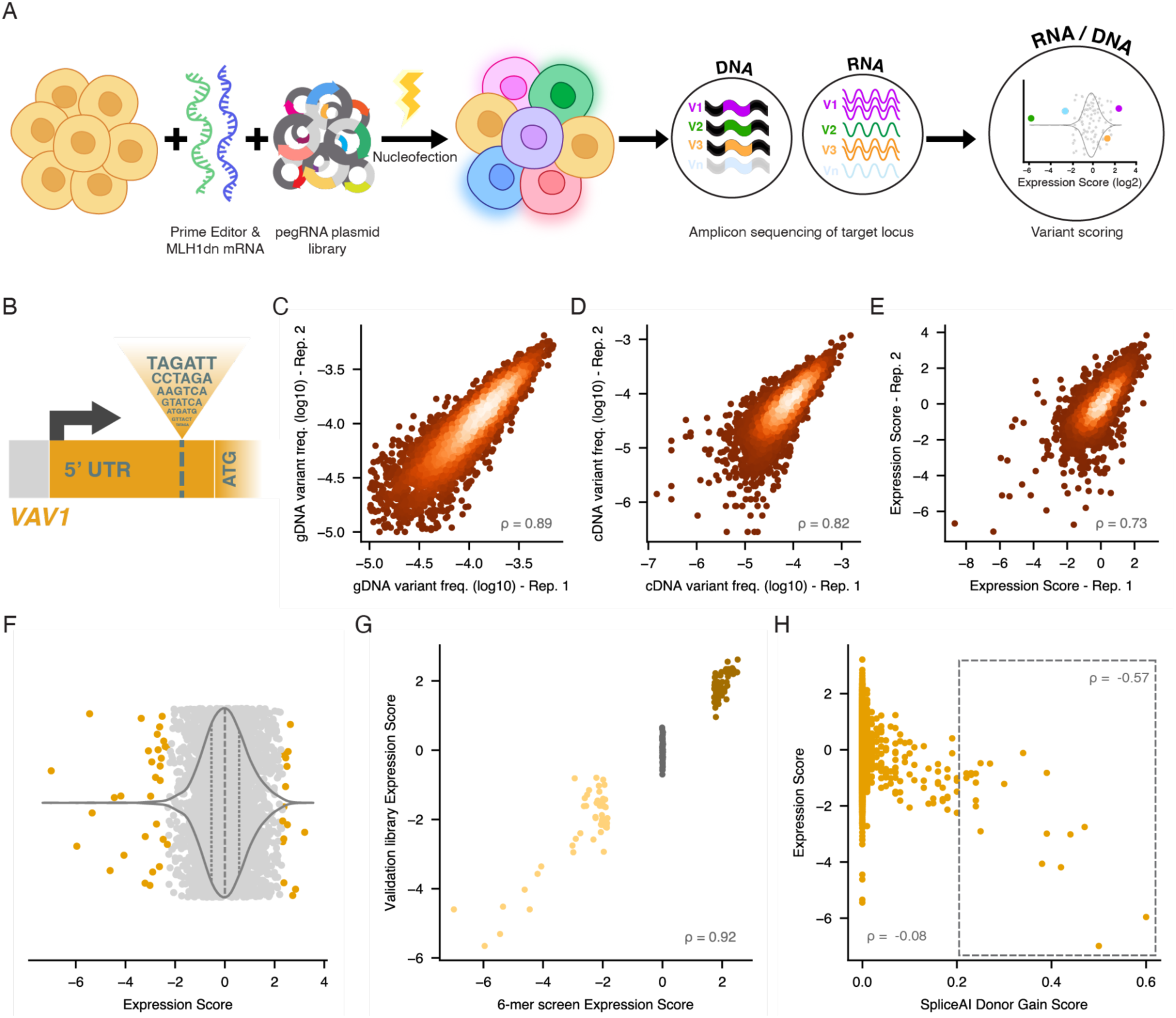
Testing effects of regulatory variants on *VAV1* expression with PETRA. **A.** Schematic of the PETRA workflow. Prime editor and MLH1dn mRNA are delivered to cells with the pegRNA plasmid library via nucleofection. Library variants are installed in transcribed regions of the target gene. Seven days post-nucleofection, amplicon sequencing of gDNA and mRNA is performed to quantify variant frequencies. An expression score is computed for each variant as the normalised log2-ratio of mRNA variant frequency over gDNA variant frequency. **B.** Schematic of the random 6-bp insertion library (*n* = 4,096) targeting the 5’ UTR of *VAV1*. **C.** Correlation of gDNA variant frequencies in replicates 1 and 2 for *n* = 3,902 insertions post-filtering (Spearman’s ⍴). Correlations across all replicates are in Supplementary Figure 1C. **D.** Correlation of mRNA variant frequencies in replicates 1 and 2 (*n* = 3,902). **E.** Correlation of variant expression scores (log2-scaled) in replicates 1 and 2 (*n* = 3,902). **F.** Distribution of expression scores (*n* = 3,902) averaged across three biological replicates and normalised to the median. (Violin plot: dashed line, median; dotted lines, 25th and 75th percentiles, all points shown with orange indicate *p* < 0.01 under a normal distribution. **G.** Correlation of expression scores for *n* = 150 selected insertions from the *VAV1* 6-mer screen re-tested using an independent validation library. (Dark brown, high-scoring in initial screen; grey, neutral-scoring; yellow, low-scoring.) **H.** Expression scores versus SpliceAI donor gain scores for *n* = 3,902 insertions. Spearman’s ⍴ is shown for all insertions and for only insertions with SpliceAI scores over 0.20 (dotted box).

To demonstrate PETRA in Jurkat T cells, we first analysed effects of variants on *VAV1* expression. Upregulation of *VAV1* using CRISPRa has been shown to enhance T cell effector function in multiple assays^24^. Here, we tested whether increased *VAV1* mRNA could be achieved by introducing edits endogenously. To this end, we cloned a library of pegRNAs designed to install every possible 6-bp insertion (*n* = 4,096) at a single site in the gene’s 5’ untranslated region (5’ UTR) immediately downstream of the *VAV1* transcription start site (TSS; **Figure 1B**, **Supplementary Figure 1A**). A pegRNA architecture was selected on the basis of predicted editing efficiency^25^ (see **Methods**). The pegRNA library was delivered to Jurkats via nucleofection with mRNAs encoding PE7^26^ and MLH1dn^27^. Seven days post nucleofection, cells were harvested and the target locus was amplified and sequenced from genomic DNA (gDNA) and mRNA. For each variant, an expression score was computed as the log2-ratio of variant frequency in mRNA over variant frequency in gDNA, normalised to the median.

In this experiment, 6-bp insertions were installed with a combined editing efficiency of 50.4% (**Supplementary Figure 1B**), allowing successful scoring of 3,902 variants (95.3% of the library) after removal of variants seen at low gDNA frequencies (less than 1.0 × 10^−5^) (**Supplementary Table 1**). Across three replicate transfections, variant frequencies in gDNA and mRNA and variant expression scores were highly correlated (**Figure 1C-E**, **Supplementary Figure 1C**). We defined 46 insertions with significant effects on *VAV1* expression (*p* < 0.01), including insertions that decreased expression to less than 20% of baseline and insertions that increased expression by more than 5.0-fold (**Figure 1F**). A subset of 150 variants that scored either highly, lowly, or neutrally were re-tested in an independent library (**Supplementary Table 2**). Expression scores from this validation experiment were strongly correlated to those from the larger experiment (Spearman’s ρ = 0.92; **Figure 1G, Supplementary Figure 1D**), confirming PETRA’s reproducibility.

Variants introduced to the 5’ UTR may impact mRNA levels through a variety of mechanisms, including sequence-specific effects on transcription initiation, transcript stability, and pre-mRNA splicing. Reasoning that variants disruptive of *VAV1* splicing may be underrepresented in mRNA, we scored insertions for their predicted splice effects using SpliceAI^3^ (**Figure 1H; Supplementary Table 1**). While most insertions had no predicted impact on splicing, several variants predicted to create new donor splice sites had low expression scores, consistent with aberrant splicing leading to variant depletion in mRNA. Considering only insertions with SpliceAI scores greater than 0.20, we observed a correlation between SpliceAI scores and expression scores of ρ = −0.57, confirming expression scores reflect biological effects of variants on mRNA levels.

### Scoring 13,935 variants in total across four genes

We next extended PETRA to three additional genes: *IL2RA*, *OTUD7B* and *CD28*. Increased expression of each of these genes has been reported to enhance T cell effector function^24,28^. We designed PETRA libraries once more comprising all possible 6-bp sequences to be inserted at a single site downstream of each gene’s TSS and achieved insertion efficiencies between 31.3% and 70.3% (**Supplementary Figure 1B, E-G**). As for *VAV1*, we removed insertions not installed efficiently and assigned expression scores to the remaining 10,033 variants, increasing the total number of variants scored across the four genes to 13,935 (85.1% of library variants; **Figure 2A, Supplementary Table 1**). Once more, expression scores were correlated across replicates **(Supplementary Figure 2A**), and independently assayed subsets of variants validated highly and lowly scoring insertions from the larger experiments (**Supplementary Figure 2B-E, Supplementary Table 2**). Concordance in the validation experiment was weakest for *IL2RA*. This result is consistent with *IL2RA* transcript levels being low in Jurkats at baseline (**Supplementary Figure 2F**), and not inefficient prime editing, as *IL2RA* insertions were installed with high efficiency (**Supplementary Figure 1B**).

**Figure 2.**
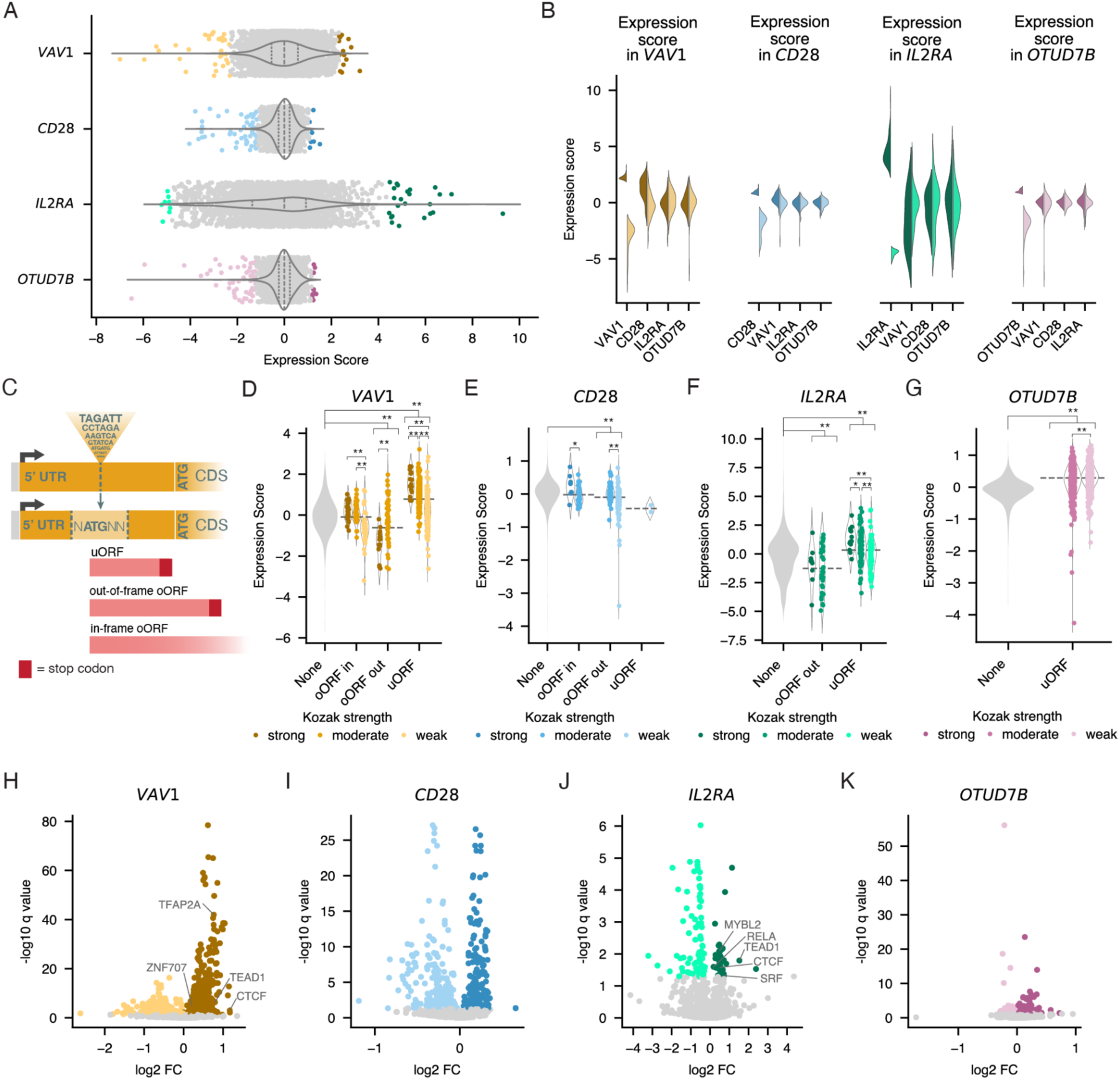
Exploring expression effects of 13,935 non-coding variants across four genes. **A.** Distribution of expression scores for 6-bp insertions targeting the 5’ UTR of *VAV1* (*n* = 3,902), *CD28* (*n* = 3,930), *IL2RA* (*n* = 2,419), and *OTUD7B* (*n* = 3,684), averaged across biological replicates (*n* = 3 for *VAV1* and *CD28*; *n* = 2 for *OTUD7B* and *IL2RA*) and median-normalised. (Violin plot: dashed line, median; dotted lines, 25th and 75th percentiles, coloured points represent standard normal distribution test *p* < 0.01). **B.** For the top 50 (dark) and bottom 50 (light) insertions scored at each locus (x-axis), the split violin plots show expression scores of the same insertion sequences introduced to each gene (panels). Insertion sequences are defined in the 5’ to 3’ orientation of the sense strand. **C.** Schematic of new ORF classification. uORF = upstream ORF, oORF = overlapping ORF. **D-G.** Violin plots of expression scores per gene, grouping variants by ORF type and Kozak sequence strength. Horizontal dotted lines indicate median by ORF type. (* = *q* < 0.05, ** = *q* < 0.01 for Mann-Whitney U-test with BH correction; oORF in = in-frame oORF, oORF out = out-of-frame oORF) **H-K.** For each TF expressed in Jurkats, a log2-fold-change (log2FC) was calculated as the difference in median expression score between insertions creating a FIMO-predicted binding site for the TF and other insertions at the site. Volcano plots show the log2FC for each TF motif with q-values from a two-sample Kolmogorov-Smirnov test with BH correction. Colours indicate *q* < 0.05 for TF motifs associated with increased (dark colours) or decreased (light colours) expression. TF motifs used to design insertions in subsequent experiments are labelled.

Across the four genes, 204 variants significantly modulated expression (*p* < 0.01), with 57 increasing expression and 147 decreasing expression (**Figure 2A**). Exploring whether the same 6-bp insertion sequences influence expression similarly across loci, we found most effects to be highly context-dependent. For instance, the 50 highest and lowest scoring 6-bp sequences inserted at each site tended to score neutrally at each of the other loci (**Figure 2B**). This high degree of locus specificity may be attributable to differences in adjacent sequence context or interactions with nearby regulatory elements.

### Exploring mechanisms by which sequences impact mRNA levels

Towards explaining sequence-specific effects on expression, we first examined splicing. Splicing effects were predicted for insertions at each locus using SpliceAI (**Supplementary Figure 2G-J**). As for *VAV1*, higher SpliceAI scores predicted lower expression scores for insertions in *CD28*. Several variants strongly predicted to impact *OTUD7B* splicing had very low expression scores, but overall the correlation was insignificant (ρ = −0.02, *p* = 0.2). No *IL2RA* insertions had SpliceAI scores above 0.04. Across all four genes, the majority of significantly scored variants had SpliceAI scores less than 0.04, suggesting other mechanisms explain their effects.

New open reading frames (ORFs) created in the 5’ UTR can lower protein expression by impacting ribosome loading and translation, yet how new ORFs impact mRNA levels is less clear^29,30^. To explore this, all 2,031 insertions that created new 5’-ATG sequences were classified by Kozak sequence strength and the type of ORF created. Upstream ORFs (uORFs) are ORFs that end upstream of the original start codon, whereas overlapping ORFs (oORFs) extend into the original ORF and can be either in-frame or out-of-frame (**Figure 2C**). Excluding variants with SpliceAI scores greater than 0.1, variants creating uORFs had significantly higher expression scores than variants not creating new ORFs across loci (Mann-Whitney U test with Benjamini-Hochberg (BH) correction for *VAV1*: *q* = 2.50×10^−45^; *IL2RA*: *q* = 2.79×10^−4^; *OTUD7B*: *q* = 2.18×10^−147^), with the exception of *CD28*, where only three uORF variants were tested (**Figure 2D-G**; **Supplementary Table 3**). This effect was dependent on Kozak sequence strength, with insertions creating strong Kozak sequences in *VAV1* and *IL2RA* scoring highest. Meanwhile, insertions creating out-of-frame oORFs scored more lowly than insertions not introducing 5’-ATG sequences (Mann-Whitney U test with BH correction for *VAV1*: *q* = 3.10×10^−5^; *CD28*: *q* = 2.19×10^−6^; *IL2RA*: *q* = 1.23×10^−3^). No difference was observed for variants creating in-frame oORFs, indicating the negative effect for out-of-frame oORFs is likely attributable to nonsense-mediated decay. While this analysis shows that new ORFs can influence mRNA expression, only 38 of 204 (18.6%) variants scored as hits created new ORFs.

We next characterised how the creation or disruption of transcription factor binding sites (TFBSs) may impact expression. For this analysis, we predicted the binding of each transcription factor (TF) expressed in Jurkats^31^ to each variant allele using FIMO^32^ and the JASPAR database^33^, once more excluding variants with SpliceAI scores greater than 0.10. Across loci, insertions were not predicted to disrupt pre-existing TFBSs (**Supplementary Figure 1A,E-G**).

To examine effects of newly created TFBSs, we asked whether expression scores differed for variants predicted to create a binding site for a given factor compared to all other variants at the site. This revealed 555 motifs associated with increased expression of at least one gene and 567 motifs associated with decreased expression (**Figure 2H-K, Supplementary Table 4**). Interestingly, no motif had a consistently significant effect across the four genes, and very few motifs were consistent across three genes (**Supplementary Figure 2K-L**).

### Single and combinatorial delivery of TFBSs to increase expression

We next set out to explore positional and combinatorial effects of insertions, with the goal of inducing larger changes in expression. For this purpose, we designed pegRNAs to introduce the MYBL2 motif to each of three loci in the *IL2RA* 5’ UTR, as 6-bp sequences predicted to create MYBL2 binding sites were associated with increased *IL2RA* expression (**Figure 2J**). We also designed three additional pegRNAs to insert the EGR1 motif at the same loci, which was not associated with altered *IL2RA* expression. PETRA was then performed using a pool of these six pegRNAs, allowing a set of 27 alleles to be generated and scored in a single transfection (**Figure 3A**).

**Figure 3.**
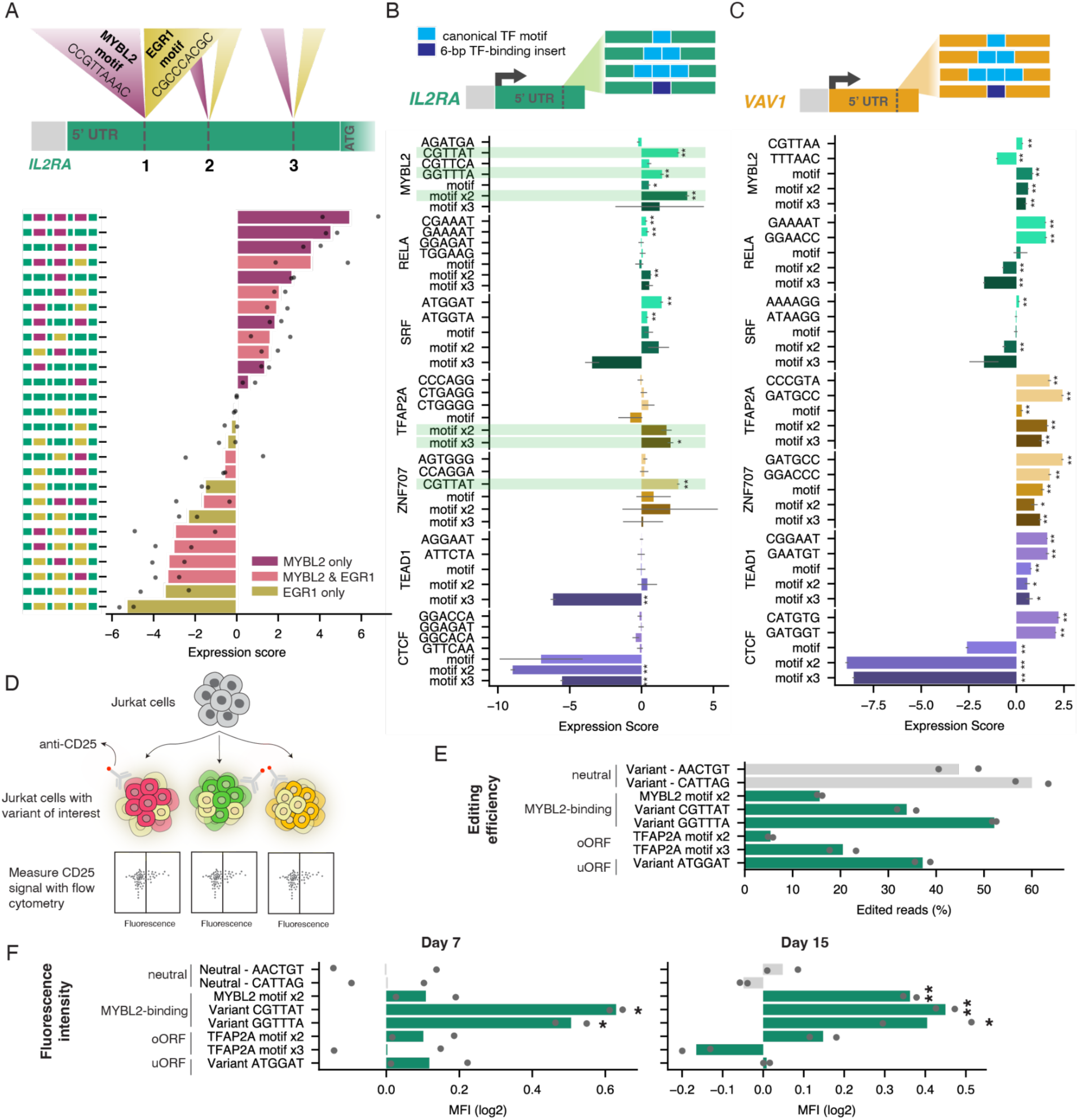
TF motifs identified with PETRA modulate RNA and protein expression. **A.** Schematic shows insertions of either the MYBL2 or EGR1 motif at each of three sites in the 5’ UTR of *IL2RA*. Six pegRNAs programming the insertions were co-delivered to cells to achieve combinatorial editing. The barplot shows PETRA expression scores for each of 27 alleles observed following editing, normalised to unedited (insertions depicted at left). Points represent independent replicates and bars represent the average across replicates. **B-C.** Schematics show pegRNA libraries designed for *IL2RA* (B) and *VAV1* (C) to deliver select TF motifs individually or in tandem to a single site within each gene. High-scoring 6-bp sequences predicted to bind each TF were included, as well as controls. Expression scores, normalised to control insertions, are plotted below, grouped by TF motif. Error bars represent standard error across three replicates. Colors indicate the reason for TF motif inclusion (green shades: hit in *IL2RA* 6-bp screen; yellow shades: hit in *VAV1* 6-bp screen; purple shades: hit in both screens. Variants individually tested for protein expression are highlighted in green. Six-bp insertions predicted to create multiple TF motifs are plotted under each corresponding motif. (** = *q* < 0.01 and * = *q* < 0.05, Welsch’s unequal variances two-sample t-test against neutral variants with BH correction) **D.** Schematic for assessing variant effects on protein expression via flow cytometry. Delivery of individual pegRNAs resulted in mixed populations of cells with one or more edited allele or no edits. **E.** Average editing efficiency of insertions tested individually in Jurkats, with dots indicating individual replicates (light grey: controls; green: high expression scores). **F.** Cell populations transfected with individual pegRNAs were activated with PMA for 24 h, anti-CD25 stained, and assessed by flow cytometry at day 7 (left) and day 15 (right) post-editing. Barplots represent mean fluorescence intensity (MFI) for CD25 normalised to neutral variants. (** = *q* < 0.01, * = *q* < 0.05, two-sample Student’s t-test versus neutral controls with BH correction)

MYBL2 binding sites individually increased expression to varying degrees (**Figure 3A, Supplementary Table 5**). Insertion at site 2 led to the highest increase among single inserts (expression score = 2.69), followed by insertion at site 1 (expression score = 1.37), validating the motif’s predicted effect. Intriguingly, combinations of MYBL2 insertions showed largely additive effects on log2-scaled expression scores. For example, the expression score of the MYBL2 motif double insertion at sites 1 and 2 was 4.57, and the score of the MYBL2 motif triple-insertion was 5.48, corresponding to a 44-fold increase in mRNA expression compared to unedited alleles. Meanwhile, single EGR1 insertions either had little effect or decreased expression, while multiple EGR1 insertions more strongly reduced expression. For example, the EGR1 triple insertion led to a 40-fold reduction in mRNA (**Figure 3A**). The additive nature of expression effects was corroborated by introducing sets of MYBL2 or EGR1 motifs individually (**Supplementary Figure 3A-B**).

We next explored whether multiple TF motifs inserted via single pegRNAs could be leveraged to increase expression. We selected the MYBL2, SRF, and RELA motifs associated with increased *IL2RA* expression and the TFAP2A and ZNF707 motifs associated with increased *VAV1* expression (**Figure 2H,J**). We also selected the TEAD1 and CTCF motifs, which were associated with higher expression of both genes. We then designed pegRNA libraries targeting the same sites in *IL2RA* and *VAV1* to which random 6-bp sequences were inserted, this time delivering up to three copies of each motif in tandem (**Figure 3B-C**). For each locus, we also included 6-bp insertions predicted to bind each TF that had scored highly before, as well as additional edits comprising positive and negative controls. Final libraries included 59 pegRNAs for *VAV1* and 77 pegRNAs for *IL2RA*. Editing rates were sufficient to score all variants, though larger insertions were created less frequently, consistent with previous studies^34^ (**Supplementary Figure 3C-J**).

Expression scores revealed that most but not all TF motif insertions increased expression of the genes for which they were selected (**Figure 3B-C; Supplementary Tables 6-7**). The highest scoring insertion at the *IL2RA* locus was the double insertion of the MYBL2 motif, which increased expression over 9-fold (**Figure 3B**). For the RELA and SRF motifs, *IL2RA* expression was significantly increased by the 6-bp insertions identified in screening, but not by the designed motif insertions, with the exception of double RELA motif insertion, which increased expression 1.5-fold. All nine TFAP2A, ZNF707, and TEAD1 motif insertions increased *VAV1* expression (**Figure 3C**), but not to a greater extent than the top-scoring 6-bp sequences for each motif. While TEAD1 motif-containing sequences failed to increase *IL2RA* expression, triple insertion of the TFAP2A motif (identified as a hit in the *VAV1* screen) led to more than a 3-fold increase. In contrast to 6-bp insertions predicted to bind CTCF, insertion of either one or multiple 13-bp CTCF motifs strongly decreased expression at both loci. We repeated the experiment for *IL2RA* in cells activated with 100 ng/ml PMA and found expression scores to correlate reasonably well to those from resting Jurkats (**Supplementary Figure 3K**). In summary, these experiments demonstrate that delivery of multiple TF motifs via single pegRNAs can drive large changes in RNA expression.

### Testing effects of insertions on protein expression

Variants that alter mRNA expression may not necessarily alter protein expression. Therefore, we individually assessed a panel of six variants with high *IL2RA* expression scores for effects on CD25 levels using flow cytometry. Specifically, we chose three variants predicted to introduce MYBL2 binding sites as well as three variants that create new ORFs and thus may interfere with translation. We also included two 6-bp insertions that scored neutrally in our large screen as controls.

We delivered pegRNAs for each insertion to cells individually, and measured CD25 expression by flow cytometry on days 7 and 15 after activating cells for 24 hours with PMA (**Figure 3D**). Editing efficiencies, determined via amplicon sequencing on day 7, were less than 60% for all samples (**Figure 3E**), meaning both edited and unedited alleles contributed to total CD25 expression in each cell population. Nonetheless, we observed increased mean fluorescence intensities for each of the three samples to which sequences predicted to bind MYBL2 were introduced (**Figure 3F, Supplementary Table 8**). The more efficiently installed 6-bp insertions significantly increased expression at both day 7 and day 15, whereas the double MYBL2 motif insertion showed a significant increase only on day 15. In contrast, the three insertions that create new ORFs did not significantly change CD25 protein expression despite having scored highly for mRNA expression (**Figure 3F**). The same trends were observed for the percentage of CD25^+^ cells in each sample (**Supplementary Figure 3L**), confirming PETRA enables identification of edits that increase expression of target proteins.

### PETRA in primary human T cells

For potential therapeutic applications, it may be valuable to identify edits with desired effects on expression in primary human cells. Towards this goal, we performed PETRA in human CD3+ T cells obtained from healthy donors (**Figure 4A**). CD3+ T cells were activated with anti-CD3, anti-CD28, and anti-CD2 antibodies on day 0 and day 14 following isolation. Three days post re-activation, cells were nucleofected with mRNA encoding PEMax^27^ and MLH1dn, as well as the pegRNA library to install all possible 6-bp insertions in the *IL2RA* 5’ UTR. On day 7, average editing was 12.3% across biological replicates, allowing expression scores to be derived for 3,143 insertions (**Supplementary Figure 4A-C, Supplementary Table 9**).

**Figure 4.**
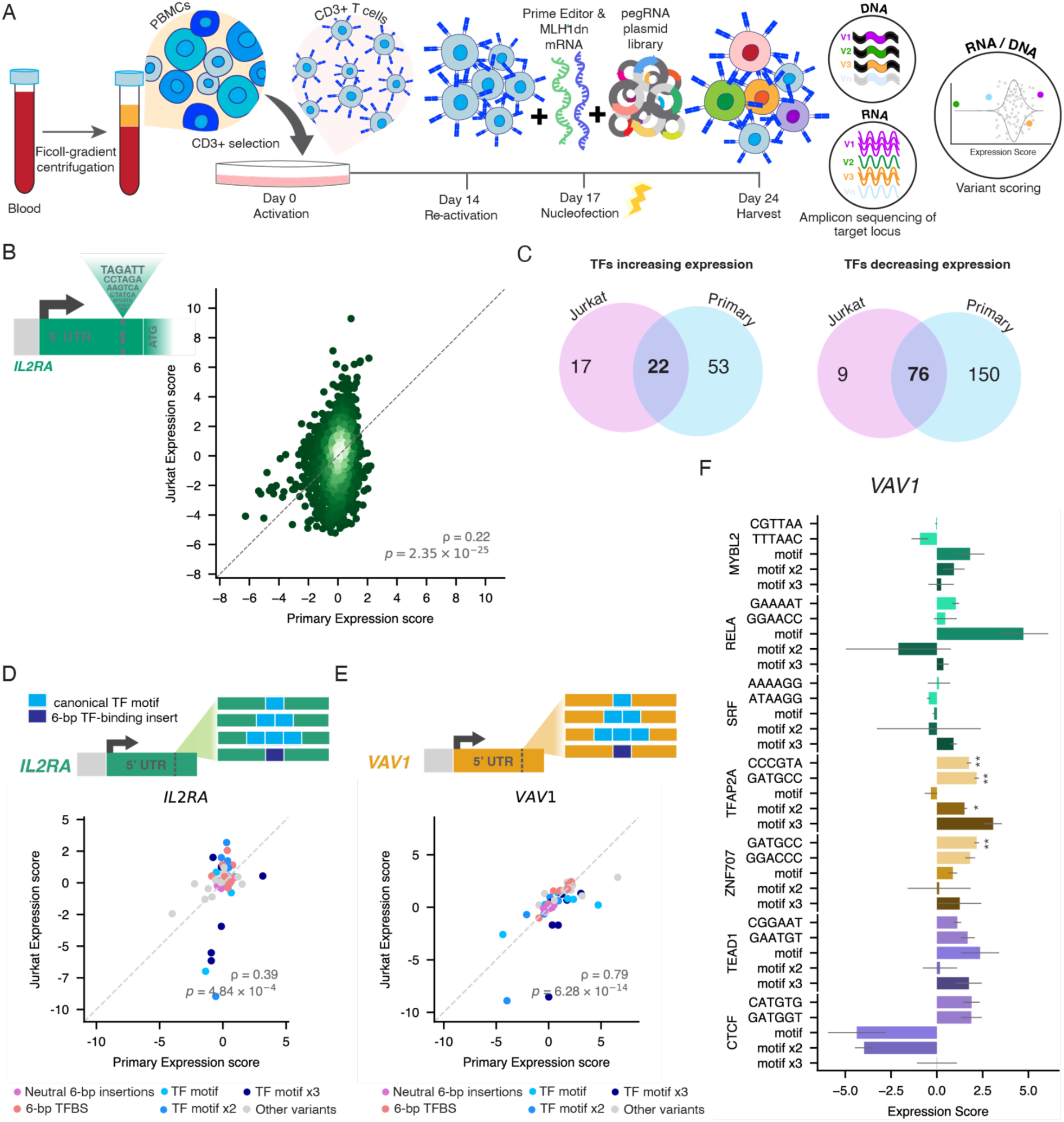
Testing effects of regulatory variants in human primary T cells with PETRA. **A.** CD3+ primary T cells selected from donor PBMCs are activated with anti-CD2, anti-CD3, and anti-CD28 on day 0 and re-activated on day 14. Three days after re-activation, prime editing is performed to deliver a library of variants scored for effects on mRNA expression. **B.** Correlation (Spearman’s ρ) of expression scores for *n* = 2,269 *IL2RA* 6-bp insertions assayed in Jurkat and primary human T cells. **C.** Venn diagrams showing the overlap of TF motifs associated with increased or decreased *IL2RA* expression scores in Jurkat and primary T cells. **D-E.** The same pegRNA libraries designed to deliver TF motifs individually or in tandem to Jurkat cells (Figure 3B-C) were assayed in primary T cells. Correlations (Spearman’s ρ) between expression scores from Jurkat and primary T cells are plotted for *n* = 77 *IL2RA* variants (D) and *n* = 59 *VAV1* variants (E), coloured by insertion type. **F.** Expression scores, normalised to neutral insertions, are plotted for *VAV1* TF motif insertions assayed in CD3+ primary human T cells. Error bars represent standard error across three biological replicates. ** = *q* < 0.01 and * = *q* < 0.05, Welsch’s unequal variances two-sample t-test against neutral variants with BH correction.

Expression scores from primary T cells were only modestly correlated to Jurkat-derived expression scores (ρ = 0.22, *p* = 2.35 × 10^−25^; **Figure 4B**), yet TF motif analysis identified many motifs associated with increased expression in both primary T cells and Jurkats, including MYBL2, RELA, TEAD1, and CTCF (**Figure 4C, Supplementary Table 10**). As in Jurkats, uORFs were associated with higher expression scores, whereas out-of-frame oORFs were associated with lower expression scores (**Supplementary Figure 4D**).

As several TF motifs were associated with increased *IL2RA* expression in both Jurkats and primary cells, we additionally assayed the smaller pegRNA libraries designed to install specific TF motifs at *IL2RA* and *VAV1* in primary T cells. For *IL2RA*, expression scores were modestly correlated between Jurkats and primary T cells (ρ = 0.39, *p* = 4.84 × 10^−4^; **Figure 4D, Supplementary Figure 4E; Supplementary Table 11**). The correlation was similar when using Jurkat expression scores from cells activated with PMA (ρ = 0.36, *p* = 1.11×10^−3^; **Supplementary Figure 4F**). Interestingly, *VAV1* expression scores were much more highly correlated between cell types (ρ = 0.79, *p* = 6.28 × 10^−14^; **Figure 4E; Supplementary Table 12**), with many of the same insertions that increased expression in Jurkats also scoring positively in primary T cells (**Figure 4F**). Collectively, these results establish the feasibility of PETRA in primary T cells.

### Comparing PETRA data to outputs from predictive models of expression

Models for predicting RNA expression from DNA sequence have improved, yet how well they perform at predicting expression effects of variants introduced by editing remains uncertain^7,35^. We assessed three recent artificial intelligence (AI) models, Enformer^5^, Borzoi^36^ and AlphaGenome^37^, by comparing their cell-type specific predictions for cap analysis of gene expression (CAGE) and RNA-seq to PETRA expression scores (**Supplementary Tables 13-17**).

For the large libraries of random 6-bp insertions, performance varied highly across loci and output tracks (**Figure 5A**). CAGE and RNA-seq predictions were most strongly correlated with *CD28* expression scores, with a maximum correlation observed for AlphaGenome RNA-seq predictions (ρ = 0.42, *p* = 6.8 × 10^−169^). Weaker positive correlations were observed for *OTUD7B* (Enformer CAGE: ρ = 0.21, *p* = 8.4 × 10^−37^; Borzoi RNA-seq: ρ = 0.13, *p* = 1.3 × 10^−14^; AlphaGenome RNA-seq: ρ *=* 0.17, *p =* 9.2 × 10^−26^) and *VAV1* (Enformer CAGE: ρ = 0.17, *p* = 7.5 × 10^−26^; Borzoi RNA-seq: ρ *=* 0.01, *p* = 5.1 × 10^−1^; AlphaGenome RNA-seq: ρ *=* 0.15, *p* = 3.8 × 10^−22^). Meanwhile, predictions for *IL2RA* from all three models were anti-correlated with expression scores from both Jurkat and primary T cells (**Figure 5A**).

**Figure 5.**
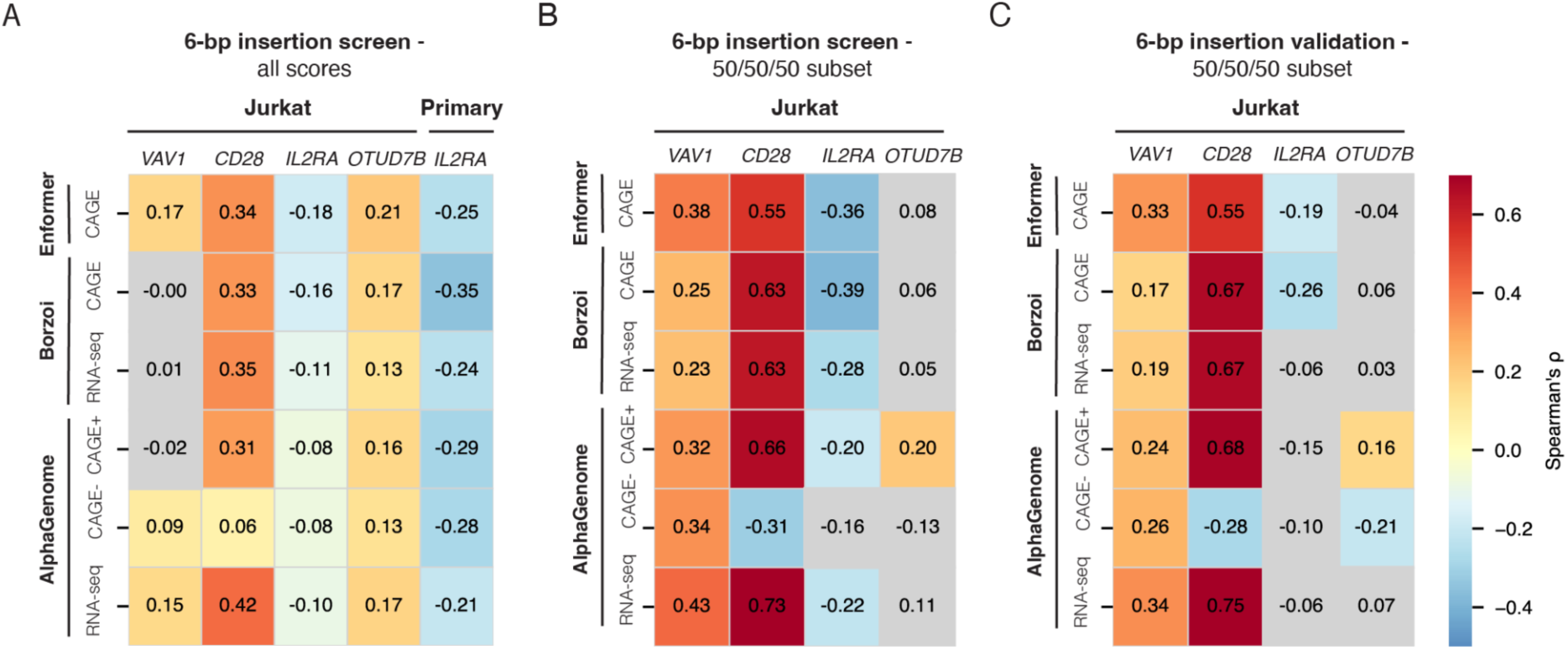
Comparing expression scores to predictions from sequence-to-expression models. **A.** Spearman correlations between expression scores for random 6-bp insertions and RNA-seq or CAGE predictions, considering all insertions scored in 6-bp screening experiments (Jurkat: *VAV1 n* = 3,902, *CD28 n* = 3,930, *IL2RA n* = 2,419, and *OTUD7B n* = 3,684; primary T cells: *IL2RA n* = 3,143). **B.** Spearman correlations as in (A), but restricted to insertions selected for validation experiments due to having high, low, or neutral scores (*VAV1 n* = 150, *CD28 n* = 150, *IL2RA n* = 138, and *OTUD7B n* = 150). **C.** Spearman correlations for the same insertions as in (B), except using expression scores from validation experiments (*VAV1 n* = 150, *CD28 n* = 150, *IL2RA n* = 138, and *OTUD7B n* = 150). For each analysis, expression scores from Jurkats were compared to Jurkat-specific predictions, while those from primary T cells were compared to primary T cell-specific predictions. Coloured (non-grey) squares indicate correlations for which *p* < 0.05.

For *CD28* and *VAV1*, correlations improved when considering only 6-bp variants re-tested on the basis of having large effects (**Figure 5B-C**), though no consistent improvement was observed for *OTUD7B* and *IL2RA*. Importantly, correlations for the same sets of variants were highly similar whether using scores from the larger screening experiments (**Figure 5B**) or the much smaller validation pools (**Figure 5C**), indicating the highly variable performance across loci is not attributable to experimental noise.

We further assessed each model’s performance against expression scores from experiments in which TFBSs were delivered to *IL2RA* and *VAV1*. Overall correlations were generally modest for alleles with MYBL2 and EGR1 binding sites installed combinatorially (**Supplementary Figure 5A-C**). It is worth noting, however, that both Borzoi and AlphaGenome were highly accurate at predicting the additive effects of MYBL2 motifs (Borzoi RNA-seq: ρ = 0.89, *p* = 6.8 × 10^−3^; AlphaGenome RNA-seq: ρ = 1.00, *p* = 4.0 × 10^−4^ for *n* = 7 alleles with MYBL2 motif insertions only). Likewise, while overall performance at predicting expression scores of TF motif insertions was modest (**Supplementary Figure 5D-O**), the low expression scores of CTCF motifs were generally well-predicted across models. Taken together, these results demonstrate how PETRA data can be used to evaluate how well computational models predict effects of edited sequences on expression, revealing highly variable performance across endogenous loci.

## DISCUSSION

We have demonstrated PETRA as a scalable means of accurately measuring how edits to genome sequence impact mRNA levels by assaying over 14,000 variants across four genes and two cell types. Further, we show that relatively large changes in expression can be engineered by 6-bp insertions identified in large-scale screening experiments, as well as by TF motifs programmed individually or in combination.

The scalability of PETRA is mediated by engineering variants in abundantly transcribed regions, facilitating sequencing of edits in mRNA and gDNA to assay effects on expression in a straightforward manner. This design enables thousands of variants per region to be queried in multiplex, and allows PETRA to be applied to multiple genes with minimal optimization. However, relying on variants to be observed in mRNA restricts the assay to specific genomic regions, which is a key limitation.

Variants in close proximity to TSSs can impact expression through multiple mechanisms. Indeed, effects on mRNA levels in PETRA experiments were associated with the creation of new splice sites, ORFs and TFBSs. These effects were largely dependent on the surrounding sequence context to which variants were introduced, explaining the high degree of locus-specificity for 6-bp insertion sequences assayed in large-scale experiments (**Figure 2B**). However, the large size of PETRA libraries facilitated identification of sequence features with consistent effects across insertion sites. For example, the MYBL2 motif was associated with enhanced expression of *IL2RA* by scoring random 6-bp insertions at a single locus. Indeed, deliberately designed MYBL2 binding sites increased expression when introduced individually or in combination to additional sites in the *IL2RA* 5’ UTR. However, MYBL2 motifs also increased expression when delivered to the *VAV1* 5’ UTR (**Figure 3C**).

While we focused on understanding how DNA sequences impact transcript levels, we also showed that select *IL2RA* variants predicted to create new TFBSs increased protein expression, as well. Notably, this was not true for the insertions we tested that created new ORFs in the *IL2RA* 5’ UTR, suggesting such sequences should be avoided if increased protein expression is desired. Though not established here, we anticipate it may be feasible to couple PETRA to readouts directly assaying protein levels or functional phenotypes of edited cells. Indeed, two recent studies in cell lines have deployed FACS-seq to link variants introduced via prime editing to protein expression changes^21,22^. These studies examined effects of variants installed upstream of each target gene, whereas PETRA identifies variants that can be inserted downstream of the TSS that impact expression. The throughput afforded by targeting the 5’ UTR allowed us to score over an order of magnitude more variants than in either previous study.

Given PETRA’s scalability and adaptability to different genomic loci, going forward we anticipate the method will be well-suited to assaying variants across many genes and cell types. It will be informative to study, for instance, how the effects of TF motifs differ by genomic position and flanking sequence context, and how the broader landscape of factor binding and chromatin state at each locus influence variant effects^38,39^. Efforts to model RNA expression from DNA sequence may benefit from PETRA data for training and benchmarking^40^. Indeed, it will be important to assess whether the highly variable performance of models across genes observed in this study can be explained by specific genomic features. Towards more robust prediction of variant effects across contexts, we anticipate that a combination of functional assays will prove ideal, for instance, combining the scalability of MPRAs with the ability to test endogenously installed variants across loci using PETRA.

For cell engineering applications, PETRA offers distinct advantages. By directly testing libraries of variants in their endogenous context, editing efficiencies and expression effects can be measured directly for loci of therapeutic relevance. We have established that PETRA identifies variants with RNA expression effects spanning a wide range. In some potential use cases, such as therapeutic editing to treat haploinsufficiency, modest expression changes that remain stable over time may prove optimal^41^.

In summary, we anticipate PETRA will substantially expand our ability to functionally test endogenous variants for effects on RNA expression across a wide range of genomic contexts and cell types. Given the scarcity of methods to scalably assay regulatory variants in their endogenous context, we anticipate PETRA will play a key role in improving our understanding of gene regulation, building improved models of variant effect, and identifying sequence changes of therapeutic benefit.

## METHODS

### Ethics

All experiments using primary human cells were approved by an internal review board of the Francis Crick Institute (approval code: 2023 FC1; 3 April, 2023), in accordance with the UK Human Tissue Act.

### Cell culture

All cells were cultured at 37°C with 5% CO2. Viability and cell concentration were determined from 10 µl of cells in media using a CellDrop Automated Cell Counter (Denovix).

#### Jurkat cells

Jurkat E6.1 cells were cultured in RPMI medium supplemented with 10% Fetal Bovine Serum (FBS) and 1% penicillin/streptomycin (Gibco 15140122). For IL2RA-targeting screens, the media was supplemented with 200 U/ml of interleukin-2 (IL-2) (Miltenyi 130-097-745). Cells were activated by supplementing the media with 100 ng/ml phorbol 12-myristate 13-acetate (PMA) (Sigma-Aldrich P1585) and 200 U/ml of IL-2 for 24 hours.

#### Primary human T cells

Human blood was obtained from the Francis Crick Institute’s Healthy Volunteer Programme from two anonymous donors. Peripheral blood mononuclear cells (PBMCs) were isolated by Ficoll (Sigma-Aldrich H8889) density centrifugation, and stored in freezing media (90% FBS + 10% DMSO) at −70°C until use. For selection of CD3+ cells, PBMCs were thawed and sorted using magnetic activated cell sorting (MACS) with anti-CD3+ beads (Miltenyi 130-097-043). The day of selection was deemed day 0. Primary T cells were cultured in X-VIVO media (Lonza BEBP02-061Q) supplemented with 5% FBS, 1% penicillin/streptomycin and 500 U/ml of IL-2. On days 0 and 14, T cells were activated using the human T Cell Activation/Expansion kit (Miltenyi 130-091-441) with beads loaded with anti-CD2, anti-CD3 and anti-CD28 antibodies. Three days after re-activation (day 17), beads were removed by magnetic separation prior to nucleofection.

### *In vitro* transcription

*In vitro* transcription (i.v.t.) was performed as described^42^ with modifications. Briefly, template plasmids including pT7-PEmax for IVT (Addgene 178113), pT7-PE7 for IVT (Addgene 214813), and pT7-MLH1dn for IVT (Addgene 178114) were amplified with primers MA001 and MA002 (all primers in **Supplementary Table 18**) using Phusion U Multiplex PCR Master Mix (ThermoFisher Scientific F564S). PCR products were purified and concentrated using AMPure XP SPRI (Beckman Coulter A63880). I.v.t was performed using HiScribe T7 mRNA Kit with CleanCap Reagent AG (New England Biolabs E2080S) with the following modifications to the manufacturer’s instructions per 20 µl reaction: addition of 0.5 µl RNase inhibitor (New England Biolabs M0314S), replacement of UTP with 0.5 µl N1-Methyl-Pseudo-UTP (New England Biolabs N0431S), reduction of GTP from 2 µl to 1 µl, use of 3 µg input DNA, and addition of 0.2 µl Yeast Inorganic Pyrophosphatase (New England Biolabs M2403S). Following 2 hours at 37°C, i.v.t reactions were incubated with DNaseI (New England Biolabs M0303L) for 15 minutes at 37°C. RNA was purified using the Monarch RNA Cleanup Kit (New England Biolabs T2050L) and quantified using Qubit RNA HS Assay Kit (ThermoFisher Scientific Q32855). Single-use RNA aliquots were stored at −70°C.

### RNA-seq

To analyse expression, 1.5 × 10^6^ unedited Jurkat cells were pelleted and resuspended in 150 µl of DNA/RNA Shield (Zymo Research R1100-50) and submitted to Plasmidsaurus’s RNA-seq service. The expression of each target gene was assessed by transcript 3’ end counting, represented as the log2-transformed counts per million plus one (CPM+1).

### PETRA experiments

#### Nucleofections and harvesting

All nucleofections were conducted using the Lonza Nucleofector II. Each nucleofection reaction included 10 µg of MLH1dn mRNA, 1 pmol of pegRNA plasmid library and 10 µg of PE7 or PEMax mRNA. A negative control nucleofected only with prime editor and MLH1dn mRNA was included in each experiment.

For Jurkat cells, 1 million cells were nucleofected per reaction using programme X-001 (Lonza VCA-1003). Two or three separate nucleofection reactions were performed per library and treated as independent biological replicates. Jurkat cells were split every three days, and 10 million cells were harvested seven days post-nucleofection.

For primary T cells, 5 million cells were nucleofected per reaction using programme T-020 (Lonza VPA-1002). Two separate nucleofection reactions were performed for the IL2RA-6-mer library and were treated as independent biological replicates. For the TF motif insertion libraries, each independent biological replicate (*n* = 2) was obtained by pooling cells from four independent nucleofection reactions. Media was replaced with fresh media three days post-nucleofection, and between 2.2 million and 4.6 million (TF motif libraries) or 5 million (*IL2RA* 6-mer library) cells were harvested seven days post-nucleofection.

Successful nucleofection was confirmed by detecting GFP fluorescence in a sample nucleofected with 2 µg of pmaxGFP plasmid (Lonza VPA-1002). Jurkat and primary T cells were cultured in antibiotic-free conditions for 72 h after nucleofection. Primary T cells were also cultured in antibiotic-free conditions for 72 h prior to nucleofection.

#### DNA and RNA extraction

Up to 10 million cells were centrifuged for 10 minutes at 350 × *g* (Jurkat cells) or 300 × *g* (primary cells). Pellets were processed for purification of DNA and RNA using the AllPrep DNA/RNA Kit (Quiagen 80204) with QIAShredder columns (Quiagen 79656). After extraction, RNA concentration was measured with the Qubit RNA HS Assay Kit (ThermoFisher Scientific Q32855) and DNA concentration was measured using the Qubit 1X dsDNA HS Assay Kit (ThermoFisher Scientific Q33231). DNA samples were stored at −20°C and RNA samples were stored at −70°C until processed for sequencing.

#### Amplicon sequencing

All PCR reactions were performed using KAPA HiFi HotStart ReadyMix (Roche 7958935001) and supplemented with SYBR Safe (Invitrogen S33102) such that the number of cycles was determined by qPCR to avoid over amplification. Products were verified by gel electrophoresis and purified and concentrated with AMPure XP SPRI Reagent (Beckman Coulter A6388). For gDNA amplification, up to 1 µg of DNA was amplified per 50 µl reaction, with up to eight reactions per sample, which were pooled prior to product purification. Reactions were complemented with an additional 2.5 mM MgCl_2_.

For RNA amplification, up to 5 µg of total RNA were used as template for a reverse transcription (RT) reaction using SuperScrip IV Reverse Transcriptase (ThermoFisher Scientific 18090050) with gene-specific primers, following the manufacturer’s instructions. Each RT product was split into ten 50 µl PCR reactions, which were pooled prior to product purification.

#### Illumina Sequencing

Amplicons containing Nextera adapters were dual-indexed and sequenced using the Illumina Nextseq 500, with 75-, 150-, or 300-cycle kits, the Illumina Miniseq with 75-cycle kits or the Illumina Novaseq 600, depending on required coverage. The length of Read 1 was sufficient to include at least 10 bp of sequence downstream from the target site, assuming successful editing of all variants at each locus.

#### Library design and cloning

pU6_pegRNA-tevopreQ1-polyT_BsmBI and pU6_pegRNA-polyT_BsmBI were prepared from pU6_pegRNA-GG-Acceptor (AddGene 132777) through an intermediate by first removing the mRFP1 cassette via PCR amplification using primers MH57 and MH58 followed by Gibson self-assembly (New England Biolabs E2621L). The resulting vector was subsequently amplified using either MH50 and MH52 or MH51 and MH52, prior to Gibson self-assembly to generate pU6_pegRNA-polyT_BsmBI and pU6_pegRNA-tevopreQ1-polyT_BsmBI, respectively.

pegRNA sequences and insertion coordinates for all libraries can be found in **Supplementary Table 19**. Plasmids pU6_pegRNA-polyT_BsmBI and pU6_pegRNA-tevopreQ1-polyT_BsmBI were linearised by BsmBI (New England Biolabs R0739L) digestion, gel-purified (Macherey-Nagel, 740609.10) and used as backbones for Gibson Assembly (New England Biolabs E2621L) to insert the pegRNA libraries. All pegRNAs used in this study included tevopreQ1 motifs^43^.

#### Random 6-bp insertion libraries

The target loci for the random 6-bp insertion libraries were chosen based on high predicted editing efficiency, inclusion in a cCRE, and avoidance of existing TF motifs. Detection of TF motifs throughout the 5’ UTR was performed by running FIMO with a detection threshold of *p* < 1 × 10^−4^. Human TF motifs were downloaded from the JASPAR 2020 database^44^ (release 8), and annotations for cCREs were retrieved from ENCODE^45^. PRIDICT^25^ was used to predict the editing efficiency of three random 6-bp insertions in every position of the 5’ UTR of each target gene. Of the pegRNAs designed to insert the three random insertions, the highest scoring design not compromising a pre-existing TF motif was used to design the pegRNA library. The 6-bp insertions used for editing prediction were replaced with a 6-bp randomer (NNNNNN), such that all pegRNAs in the library differed solely by insertion sequence. Each pegRNA design was extended at the 5′ end with an 18-bp fragment complementary to the 3′ end of the U6 promoter (U6-comp) and at the 3′ end with the tevopreQ1 sequence^43^. Oligonucleotides for each random 6-bp insertion library were synthesised as PAGE-purified Ultramers with degenerate bases (Integrated DNA Technologies).

Each library was amplified in six separate 50 µl reactions using primers MA059 and MA098 and cloned into pU6_pegRNA-polyT_BsmBI by Gibson assembly (New England Biolabs E2621L). Products were purified, concentrated via AMPure XP SPRI (Beckman Coulter A6388) clean-up, and used for transformation into Stable Competent E. Coli cells (New England Biolabs C3040H) by mixing 25 µl of cells with 5 µl of assembled plasmid and incubating on ice for 30 minutes. After a 30 second heat shock at 42°C followed by a 5-minute incubation on ice, the cells were recovered for 1 hour at 30°C with shaking, and grown overnight in liquid culture at 37°C. Plasmid libraries were extracted using ZymoPURE II Plasmid Maxiprep Kit (Zymo Research D4203) with endotoxin removal.

#### 6-bp insertion validation libraries

For the validation libraries, 50 high-scoring, 50 low-scoring and 50 neutral-scoring variants were selected from the random 6-mer insertion libraries, using the same pegRNA designs. Variants were selected based on expression scores prior to filtering, such that variants with a range of gDNA frequencies were represented. pegRNAs were ordered as part of a oligonucleotide pool (Twist Biosciences) and cloned as for the TF motif insertion libraries.

#### Combinatorial MYBL2 and EGR1 motif insertion libraries in IL2RA

Motifs for MYBL2 and EGR1 were extracted from the JASPAR 2020 database^44^ (MYBL2 motif MA0777.1: 5’-CCGTTAAAC; EGR1 motif MA0162.3-4: 5’-CGCCCACGC). PRIDICT was used to predict the efficiency of inserting both motifs in every position of the 5’ UTR of *IL2RA.* Three positions were chosen based on high predicted editing efficiency for both insertions while ensuring sufficient separation such that editing of each locus would not interfere with pegRNA spacer and primer binding site (PBS) complementarity at another target. The highest scoring pegRNA design for each insertion was chosen, and the pegRNA sequence was extended at the 5′ end with the U6-comp sequence and at the 3′ end with the tevopreQ1 sequence followed by a stuffer sequence for length uniformity and a 20-nt library-specific barcode prior to synthesis in an oligonucleotide pool (Twist Biosciences). The library-specific barcode enabled the pegRNAs for each motif (MYBL2 or EGR1) to be amplified and cloned separately, using MA104 and a library-specific reverse primer binding to the 20-nt library barcode. Following purification of each product, a second amplification was performed with MA059 and MA098. The MYBL2 motif insertion pegRNAs and the EGR1 motif insertion pegRNAs were then assembled into pU6_pegRNA-polyT_BsmBI separately, and pooled in equal quantities for experiments inserting both motifs.

#### TF motif insertion libraries

For single-motif insertions, the nucleotides from the JASPAR motif predicted to contribute most strongly to factor binding were chosen as the core motif to be inserted. For double- and triple-motif designs, tandem motifs in the same orientation were connected by adding the two nucleotides flanking the core motif. Two previously assayed 6-bp insertions predicted to bind each TF that had scored highly were also included in the library. Ten 6-bp insertions with high expression scores not predicted to bind any of the selected TFs and ten neutral insertions scoring between −0.1 and 0.1 were also included for each gene as controls, as was the 5’-GGTAAG insertion which creates a canonical splice donor variant and the four possible 1-bp insertions at each site. To create the *VAV1* TF motif insertion library, this process was performed using data from the *VAV1* random 6-mer insertion screen. To create the *IL2RA* motif insertion library, insertions were selected considering data from the *IL2RA* random 6-mer screen in both Jurkat and primary T cells. Specifically, the ten top-scoring 6-bp insertions chosen included five that scored highly in primary T cells and five that scored highly in Jurkats; neutral variants were selected as those that showed average expression scores between cell types close to zero; and an additional ten 6-bp insertions that had high expression scores in both primary and Jurkat cells (but were not necessarily among the top scoring variants in either) were also added to the library. All insertions were targeted to the same site used for the 6-bp insertion screen. For each insertion, the pegRNA design with the highest PRIDICT2^46^ score using the HEK293T model was chosen, extended with cloning adapters, and synthesised (Twist Biosciences). The pegRNA oligonucleotide libraries were then amplified as described for the combinatorial MYBL2 and EGR1 motif insertion libraries, but replacing primer MA098 with MA146. The products were then cloned into pU6_pegRNA-tevopreQ1-polyT_BsmBI.

#### Variant scoring and filtering

Single-end reads were aligned to the reference amplicon using needleall (EMBOSS v6.6.0.0) and reads with an alignment score lower than 300 were removed from analysis. Variants from the random 6-bp, validation and TF motif insertion libraries were counted by searching for exact matches to the insertion sequence plus 10 bp of flanking sequence. Variants from the combinatorial MYBL2 and EGR1 insertion libraries were counted by searching for exact matches of each possible allele, starting 10 bp upstream of insertion site 1 and ending 10 bp downstream of insertion site 3. For gDNA samples, editing efficiencies were computed as the number of reads matching an intended edit over the total number of reads.

To calculate expression scores:

1. Variant frequency in each sample was determined by dividing variant-specific reads plus a pseudocount by the total number of reads post filtering.
2. Variant frequency in RNA was divided by variant frequency in gDNA from the same sample and log2-transformed
3. The resulting ratios were normalised and averaged across replicates as follows for each library:

a. For the random 6-bp insertion libraries:

i. The log2-transformed RNA:DNA ratios were normalised to the median of all variants in the sample to produce an expression score per replicate.
ii. Expression scores per variant were averaged across biological replicates and normalised to the median average score of all variants.
iii. Variants for which the gDNA frequency was lower than a set threshold for any replicate were removed from further analysis and expression scores were normalised to the median of retained variants.
iv. To determine significant effects, expression scores were standardised to z-scores, and p-values were computed assuming a standard normal distribution without formally testing for normality. A significance threshold was set to *p* < 0.01 based on q-q plot analysis.
b. For the combinatorial MYBL2 and EGR1 motif libraries:

i. The log2-transformed RNA:DNA ratios were normalised to the RNA:DNA ratio of the unedited allele to produce an expression score per replicate.
ii. Expression scores per replicate for each variant were averaged across all independent biological replicates.
c. For the TF libraries:

i. The log2-transformed RNA:DNA ratios were normalised to the mean ratio of neutral controls to produce an expression score per replicate.
ii. To test significance, per-replicate expression scores for each variant were compared to per-replicate expression scores for all neutral variants, using Welsch’s unequal variances two-sample t-test with BH correction.
iii. Expression scores per biological replicate were averaged, and once more normalised to the mean of all neutral variants to derive final scores.

For the random 6-mer insertion libraries, variants were filtered based on frequency in gDNA. For 6-mer libraries targeting *CD28*, *VAV1*, and *OTUD7B*, the gDNA frequency threshold was set to 1 × 10^−5^. For the library targeting *IL2RA*, the gDNA threshold was set to 8 × 10^−5^. A higher gDNA threshold was used for *IL2RA* due to lower RNA expression in resting Jurkats resulting in weaker inter-replicate correlations for variant frequency in cDNA. No variants were filtered from the validation libraries, the combinatorial MYBL2 and EGR1 motif insertion libraries, or the TF motif insertion libraries tested in Jurkat and primary T cells. To assess reproducibility of editing and variant scoring, inter-replicate Spearman’s correlations for variant gDNA frequency, cDNA frequency, and expression scores were calculated.

### Analysis of sequence features in random 6-mer insertion libraries

#### uORF detection and classification

Variants leading to a new 5’-ATG (referred to as 5’-ATG variants) were classified as uORF variants if there was stop codon in-frame with the new 5’-ATG preceding the canonical ORF. Otherwise, 5’-ATG variants were classified as either an in-frame oORF variant or an out-of-frame oORF variant, depending on whether the new ORF was in-frame with the canonical ORF. The strength of the 5’-ATG context was determined based on Kozak similarity^47^. Briefly, variants were assessed for the presence of an ‘A’ or ‘G’ three nucleotides upstream of the new 5’-ATG and the presence of a ‘G’ immediately downstream. Variants were deemed “strong” if both conditions were met, “moderate” if one condition was met, and “weak” if neither condition was met.

Expression scores for variants creating different types of ORFs were compared to those of non-ATG variants using a Mann-Whitney U test with BH correction. Within each ORF type, variants with different Kozak strengths were compared the same way. Variants with SpliceAI scores over 0.1 were removed from this analysis.

#### TFBS detection

To detect potential TFBSs created by variants, FIMO^32^ was run in batch mode to scan a 46-bp window including each insertion and 20 bp of sequence on each side for human TF motifs from JASPAR 2024 (release 10)^33^. Variants were classified as predicted to bind a given TF if the TF–variant p-value was below the detection threshold parameter, which was set to 0.001. FIMO output was filtered to consider only TFs expressed in Jurkat cells (log2-transformed TPM+1 greater than 0 in batch-corrected public expression dataset 24Q4l, DepMap ID ACH-000995)^48^ and only TFBSs created by an insertion sequence. Then, for each TF, a TF motif score was calculated as the difference between the median expression score for variants predicted to bind and the median expression score for all other variants. Significance was determined using a two-sample Kolmogorov–Smirnov test with BH correction. Variants with a SpliceAI score over 0.1 were removed from this analysis.

### Prediction of variant effects with deep learning models

#### SpliceAI

To obtain predictions for variant effects on RNA splicing, SpliceAI was run from the command line, using hg38 as the reference genome, GENCODE V24 canonical files for transcript annotation, and a maximum distance parameter of 300 bp. Except where donor gain scores are specifically analysed, the maximum score among individual scores for donor gain, acceptor gain, donor loss and acceptor loss was used as a single SpliceAI score in analysis.

#### Enformer

Enformer was used to predict the effects of assayed insertions on RNA expression. For each variant, a reference and an alternative allele were created by centering the variant in a GRCh38 window with 196,608 bp on either side. Enformer predictions for 5,313 output tracks were generated for reference and alternative alleles, partitioned into 893 bins of 128 bp. Outputs across all bins were then averaged, and the predicted effect of the variant on each track was quantified as the difference between the alternative and reference alleles. Predictions were normalised using the EnformerScoreVariantsNormalized function.

Track metadata was obtained from: https://raw.githubusercontent.com/calico/basenji/master/manuscripts/cross2020/targets_human.txt. Track 4831 was used for the Jurkat-specific CAGE prediction. To obtain primary T cell-specific CAGE predictions, predictions for all primary T cell-specific tracks were averaged. A track was considered primary T cell-specific if the description contained “T cell”, “T-cell”, “Tcell”, “CD4” or “CD8” (case insensitive) with the exception of tracks 4731, 5029, 1882, 3584, 3653, 3827, and 4158.

#### Borzoi

Borzoi^36^ model and reference files were obtained from https://github.com/calico/borzoi, and params.json and model0_best.h5 were used as model parameters and weights. For each variant, alternative and reference alleles were created by centring the variant in a GRCh38 window with 262,144 bp on either side. Borzoi predictions for reference and alternative alleles were generated for 7,612 output tracks partitioned into 32-bp bins. For each bin, predictions for the forward and reverse-complement orientation were averaged. Using the gencode41_basic_nort.gtf file for gene annotations, bins were mapped to gene exons. Only bins overlapping the target gene were considered and summed to obtain a single reference and alternative prediction per track. The predicted effect of each variant on each track was then computed as the difference between the log2-transformed predictions for the alternative and reference alleles (logSED).

Track metadata was obtained from targets_human.txt. Tracks 312 and 6176 were considered as Jurkat-specific CAGE and RNA-seq predictions, respectively. Primary T cell-specific tracks were selected using the same criteria as for Enformer.

#### AlphaGenome

The AlphaGenome model (https://github.com/google-deepmind/alphagenome) was run using the AlphaGenome API. Variants were scored in batch, with organism set to “human” and sequence length set to “1 MB”. In total, 5,563 output tracks were obtained, with each track modality scored with the recommended variant scorer method. Tracks with ontology CURIE CLO:0007045 or EFO:0002796 were considered Jurkat-specific, while tracks with ontology CURIE CL:0000084 or CL:0000895 were considered primary T cell-specific. For RNA-seq predictions, only cell type-specific polyA plus RNA-seq tracks obtained with the GeneMaskLFCScorer were considered. For CAGE predictions, cell type-specific positive and negative strand predictions were considered separately. For variants with two predictions for a specific track type (corresponding to two different ontology CURIEs), a single prediction was computed by averaging the raw score of both ontology CURIEs. For each track type, AlphaGenome raw scores were used for comparison.

AlphaGenome predictions are subject to the AlphaGenome Output Terms of Use: http://deepmind.google.com/science/alphagenome/output-terms.

For all track types from all predictors, comparison to expression scores was made by computing Spearman’s and Pearson’s correlation coefficients. For analysis of *IL2RA* random 6-bp insertions assayed in Jurkat validation experiments, only insertions retained upon final filtering of the larger random 6-bp insertion screen were used. This allowed comparisons to model outputs for the same sets of insertions in Figure 5B-C.

### Quantification of variant effects on CD25 protein expression

#### pegRNA cloning

pegRNAs were selected from the TF motif insertion library targeting *IL2RA* and ordered as gene fragments (Twist Biosciences). Gene fragments were amplified and cloned into pU6_pegRNA-tevopreQ1-polyT_BsmBI by Gibson assembly (New England Biolabs E2621L). The assembled plasmids were transformed into Stable Competent E. Coli and prepared as described for pegRNA library cloning. Plasmids were sequence-verified by whole plasmid sequencing (Plasmidsaurus).

#### Editing

Plasmids expressing pegRNAs were delivered individually to Jurkat cells using the PETRA nucleofection protocol in two independent biological replicates. Each nucleofection resulted in a mixed pool of cells with a different number of edited alleles. To determine editing efficiency, 1 µg of gDNA was amplified as described for *IL2RA* gDNA samples in PETRA experiments. Amplicons containing Nextera adapters were dual-indexed and sequenced using a 150-cycle kit on the Illumina Nextseq 500, and editing efficiencies were calculated from single-end reads as described above, following alignment with needleall (EMBOSS v6.6.0.0) and removal of reads with an alignment score lower than 300.

#### IL2RA staining and flow cytometry

Jurkat cells were stained for CD25 using PE-conjugated anti-human CD25 antibody (BioLegend 302606). Cells were centrifuged, washed in PBS and incubated for 30 minutes at 4°C in the dark in staining solution (1 ml per 10^7^ cells), comprised of a 1:20 dilution of primary antibody in flow buffer (ThermoFisher Scientific 00-4222-26) complemented with 5% FBS. After staining, cells were washed twice and resuspended in flow buffer (1 ml per 10^7^ cells) for analysis. All centrifugations were performed at 350g for 10 minutes. A staining control, processed with the same protocol but without antibody in the staining buffer, was included in each experiment.

For each sample (*n* = 2 per variant), 10,000 cells were analysed in a SH800S Cell Sorter (SONY) using a 100 µm microfluidic sorting chip with the following detector and threshold settings: FSC: 4; BSC: 32%; FL1-FITC: 55%; FL2-PE: 48%. PE signal was used for detection of anti-IL2RA antibody; FITC was used to measure autofluorescence.

Flow cytometry data was analysed using FlowJo 10.10.0 software (BD Life Sciences). Raw data was filtered by gating to remove cell debris and cell doublets. The mean fluorescence intensity (MFI) from the cell population was exported for analysis. Cells with a PE signal above the maximum staining control signal were considered CD25^+^. MFI and percentage of CD25^+^ cells measurements were log2-transformed and normalised to the mean of neutral variant samples. For each variant, the MFI and percentage of CD25^+^ cells across replicates were compared to the same values obtained from neutral variants (considered as a single group) using a Student’s t-test with BH correction.

## DATA AVAILABILITY

All expression scores from PETRA experiments are available in Supplementary Tables. Fastq files from PETRA experiments will be made available prior to publication (European Nucleotide Archive accession: PRJEB102640).

## CODE AVAILABILITY

Custom scripts used to analyse PETRA experiments and to generate figures are available on GitHub: https://github.com/magdaaarmas/PETRA.

## AUTHOR CONTRIBUTIONS

M.A.R. performed all experiments and analyses. M.H. generated critical reagents and provided prime editing advice. L.C. assisted with flow cytometry and sequencing. M.A.R. and G.M.F. wrote the manuscript with input from all authors. G.M.F. supervised the work.

## ACKNOWLEDGEMENTS AND FUNDING

We thank the Crick’s Genomics Scientific Technology Platform (STP) for performing sequencing, the laboratory of James Lee for technical advice, Gita Mistry and the Human Biology STP for facilitating access to donor blood, the Flow Cytometry STP for flow cytometry assistance, and the Cell Services STP for cell line maintenance.

G.M.F. is supported by a Starting Grant from the European Research Council (Seq2Func-NC). The Francis Crick Institute receives its core funding (G.M.F.) from Cancer Research UK (CC2190), the UK Medical Research Council (CC2190), and the Wellcome Trust (CC2190). For the purpose of Open Access, the authors have applied a CC BY public copyright license to any Author Accepted Manuscript version arising from this submission.

## COMPETING INTERESTS

The authors declare no conflicts of interest.

## SUPPLEMENTARY FIGURES

**Supplementary Figure 1.**
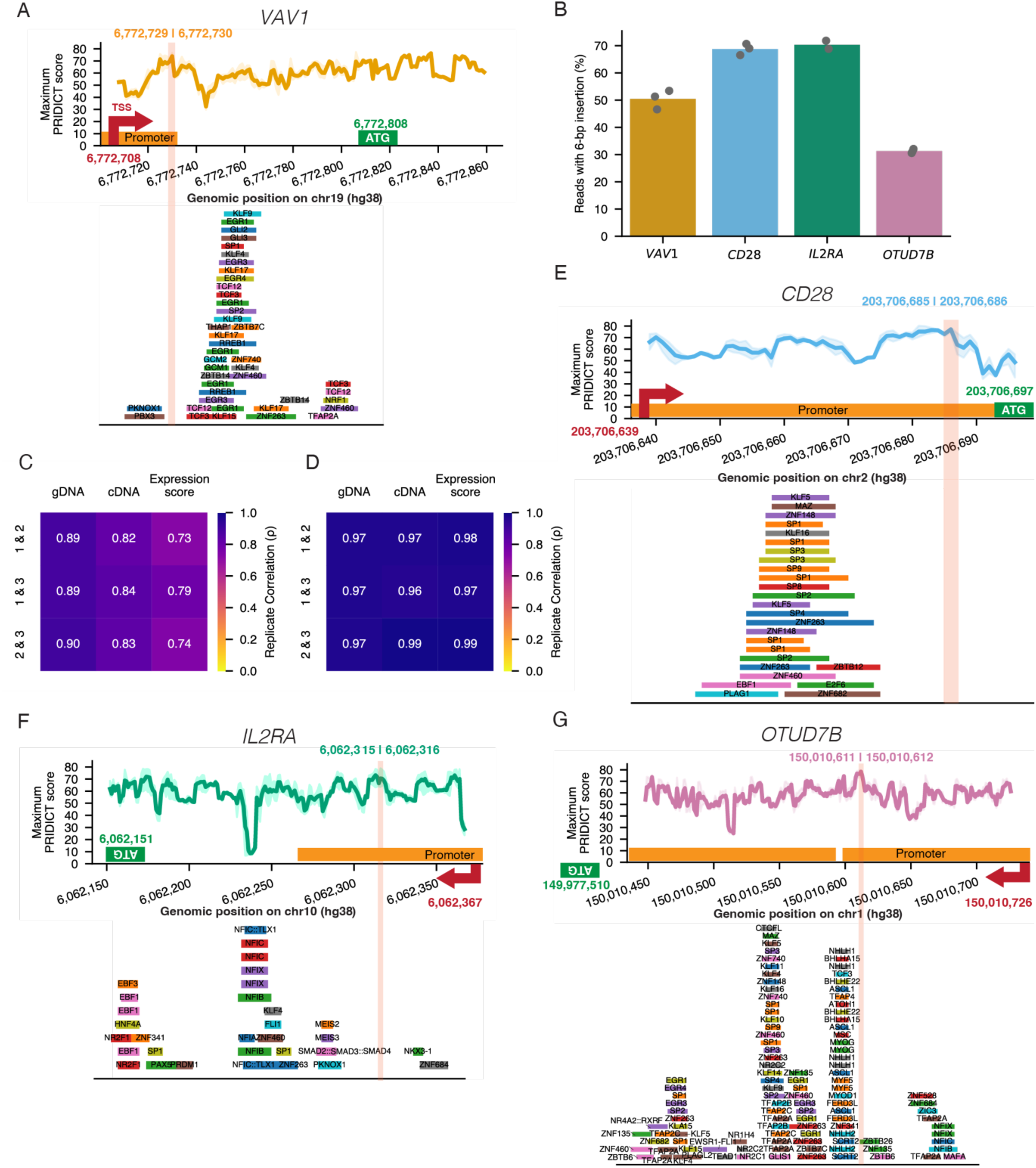
Delivery of 6-bp insertion libraries to promoter regions of four genes using prime editing. **A.** Design of 6-bp insertion library targeting the 5’ UTR of *VAV1*. The predicted editing efficiency of the top-scoring pegRNA designed to install each of three randomly selected 6-bp insertions is plotted by position. The dark line represents the average PRIDICT score of the three insertions, and the shaded area represents the range between maximum and minimum PRIDICT scores. Each gene’s ENCODE-defined promoter-like signature (PLS) candidate cis-regulatory element (cCRE) is indicated in orange. The position chosen for each 6-bp library insertion is highlighted in red, with genomic coordinates provided. Predicted TFBSs in each region are shown below. **B.** Overall 6-bp insertion efficiency measured in day 7 gDNA sequencing of each target (bar), with individual replicates as dots (*n* = 3 for *VAV1* and *CD28*, *n* = 2 for *IL2RA* and *OTUD7B*). **C.** Correlations (Spearman’s ρ) between biological replicates for variant frequencies in gDNA, cDNA and variant expression scores, for the *VAV1* 6-bp insertion library (*n* = 3,902). **D.** Correlations, as in B, between biological replicates for the *VAV1* validation library (*n* = 150). **E-G.** Design of 6-bp insertion libraries targeting the 5’ UTRs of *CD28* (E), *IL2RA* (F), and *OTUD7B* (G), as represented in (A) for *VAV1*.

**Supplementary Figure 2.**
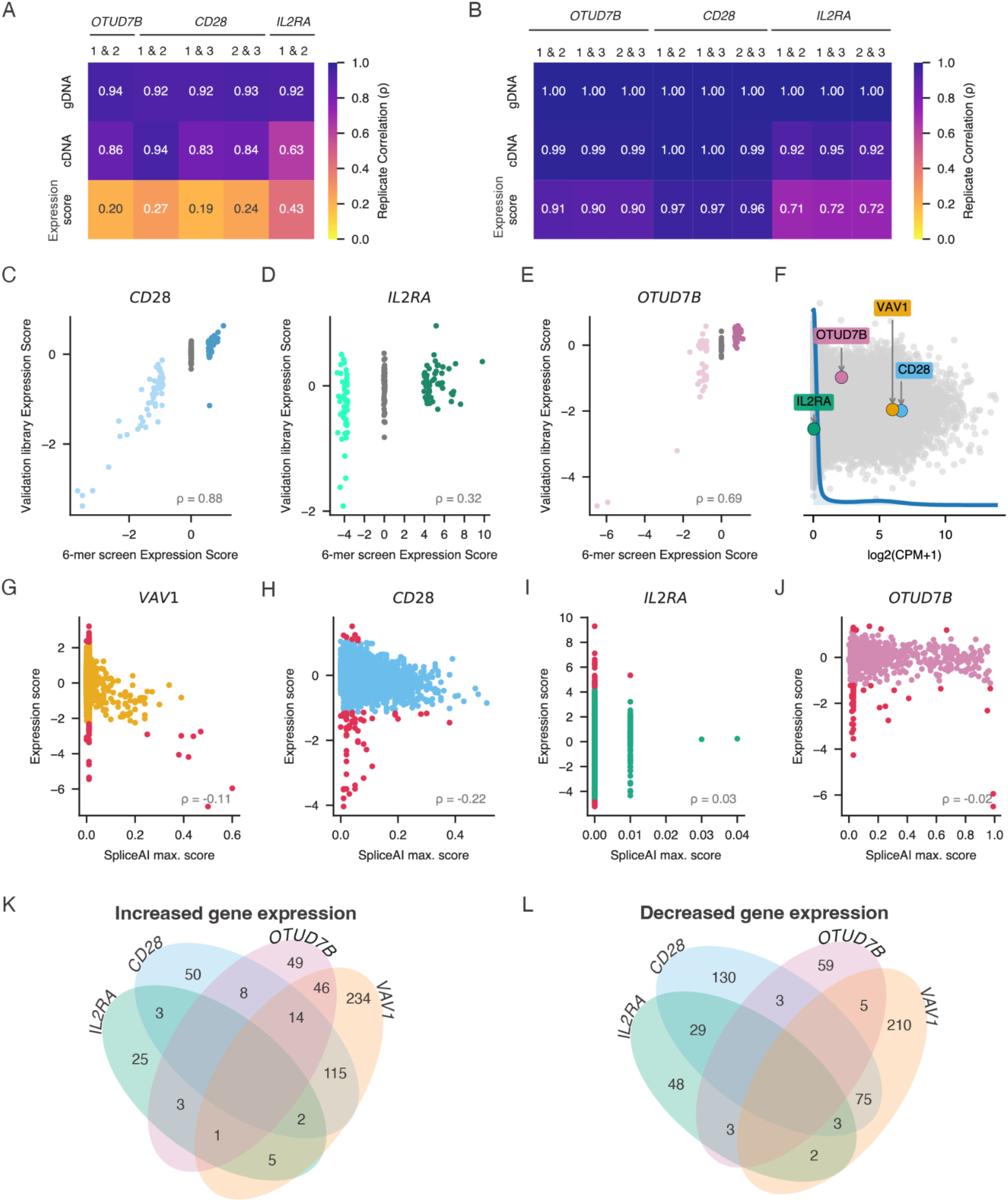
Analysis of PETRA expression scores across loci. **A.** Heatmap of Spearman’s correlations (ρ) between biological replicates for variant frequencies in gDNA, cDNA and variant expression scores, considering variants retained post-filtering on gDNA frequency. **B.** Correlations, as in A, between biological replicates for the validation libraries (*n* = 150 for each). **C-E.** Correlations (ρ) between expression scores for *n* = 150 insertions scored in the 6-mer screen and re-tested in validation libraries for *CD28* (C), *IL2RA* (D), and *OTUD7B* (E). All variants included in library design are plotted. (dark shades, high scoring; grey, neutral scoring; light shades, low-scoring) **F.** Log2-transformed counts per million (CPM)+1 values from RNA-seq of unedited Jurkat cells. **G-J.** Correlation (Spearman’s ρ) between expression scores and SpliceAI scores for each gene. Variants with significant expression scores are represented in red. (*VAV1 n* = 3,902, *CD28 n* = 3,930, *IL2RA n* = 2,419, *OTUD7B n* = 3,684) **K-L.** Venn diagrams showing the overlap of TF motifs associated with increased (K) or decreased (L) expression scores across loci.

**Supplementary Figure 3.**
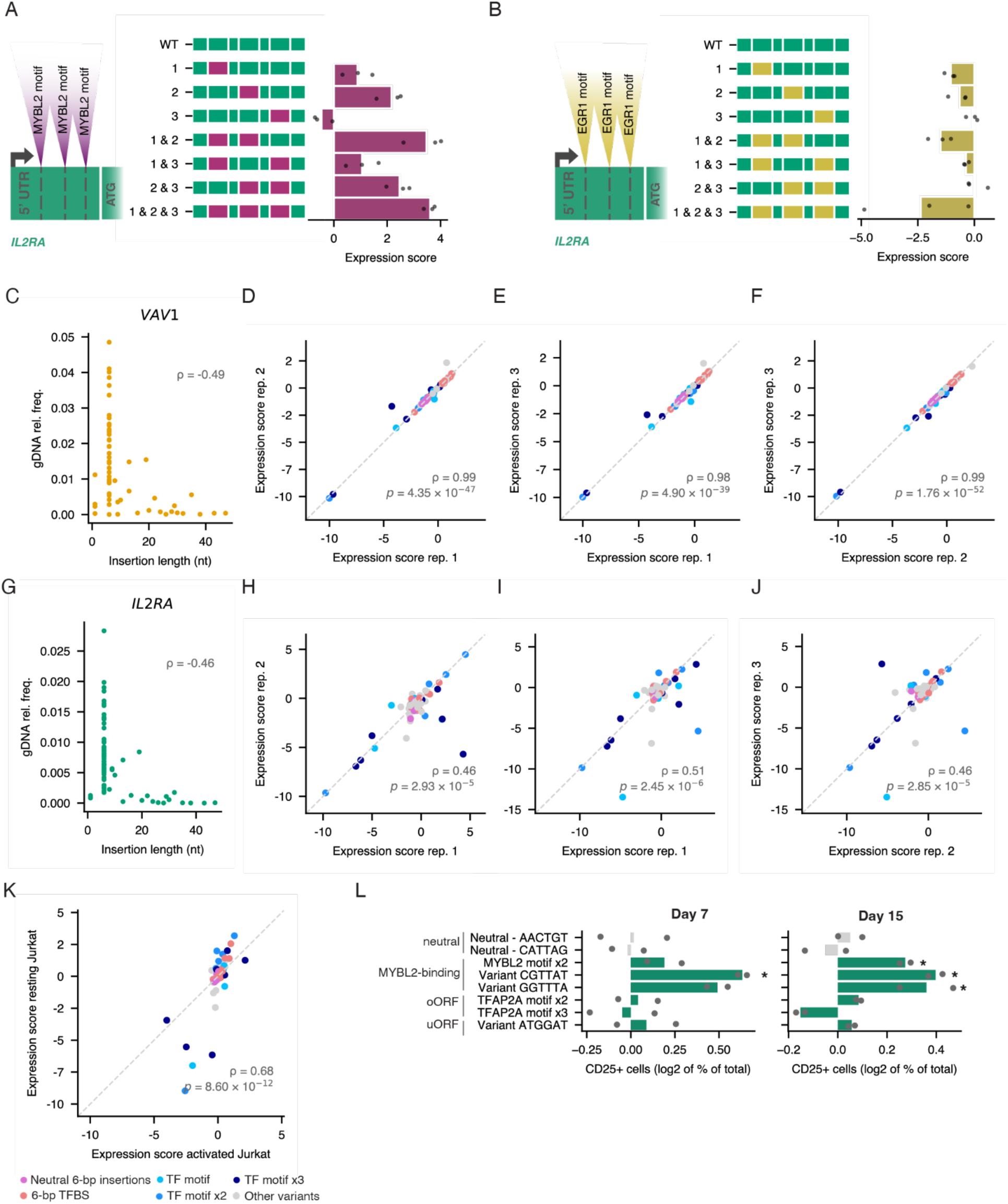
Using PETRA to assay TF motif insertions. **A-B.** Schematic: three pegRNAs designed to insert either the MYBL2 (A) or EGR1 (B) motif at each of three sites in the 5’ UTR of *IL2RA* were co-delivered to cells to achieve combinatorial editing. The barplot shows PETRA expression scores for each of 8 variant alleles observed following editing, normalised to the unedited allele (insertions depicted at left). Points represent independent replicates (n = 3) and bars represent the average across replicates. **A. C.** Correlation (Spearman’s ρ) between insertion length and gDNA variant frequency for the TF motif insertion library targeting *VAV1*. **D-F**. Scatter plots show interreplicate correlations for expression scores, with variants coloured based on insertion type (as in K). **G.** Same as C for *IL2RA*. **H-J.** Same as D-F for *IL2RA*. **K.** Correlation for expression scores for variants from the TF tandem insertion library targeting *IL2RA* between resting and activated Jurkat cells. Variants are coloured based on insertion type. **L.** Cell populations were assessed by flow cytometry at day 7 (left) and day 15 (right) after a 24 h activation with PMA. Barplots represent the frequency of CD25^+^ cells normalised to neutral variants. (** = *q* < 0.01, * = *q* < 0.05, two-sample Student’s t-test versus neutral controls with BH correction)

**Supplementary Figure 4.**
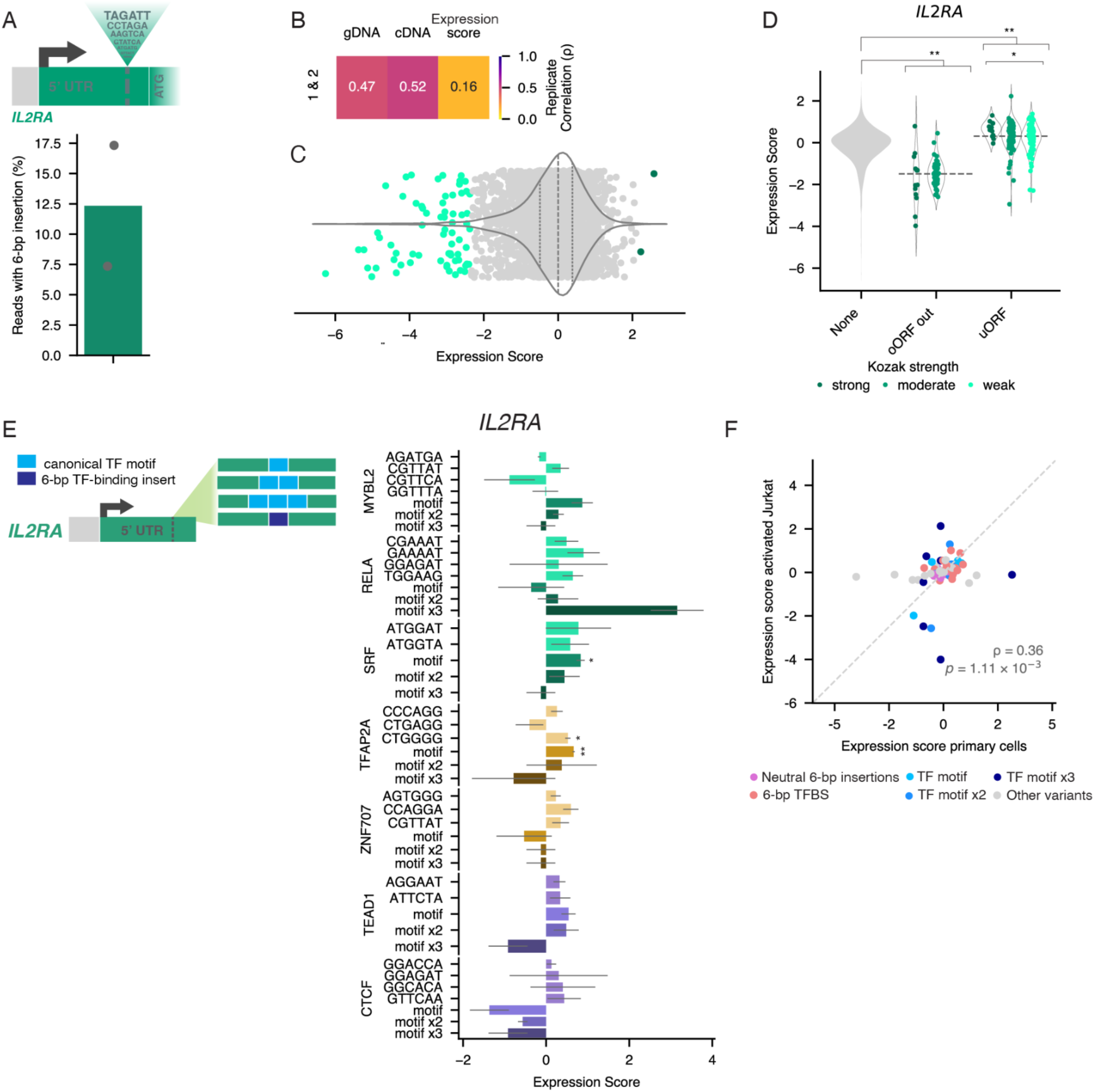
Analysis of PETRA expression scores in primary human T cells. **A.** Schematic of the random 6-bp insertion library (*n* = 4,096) targeting the 5’ UTR of *IL2RA*. Barplot shows average 6-bp insertion efficiency on day 7, with individual replicates as dots (*n* = 2). **B.** Heatmap of Spearman’s ρ between biological replicates for variant frequencies in gDNA, cDNA and variant expression scores for the 6-bp insertion library targeting *IL2RA* (*n* = 3,143 variants post-filtering). **C.** Expression scores for variants from the 6-bp insertion library targeting *IL2RA* (*n* = 3,143) averaged across two biological replicates and median-normalised. (violin plot: dashed line, median; dotted lines, 25th and 75th percentiles, green dots indicate standard normal distribution test *p* < 0.01). **D.** Violin plots of expression scores grouping insertion by ORF type and Kozak sequence strength. Horizontal dotted lines indicate median by ORF type. (* = *q* < 0.05, ** = *q* < 0.01 for Mann-Whitney U-test with BH correction; oORF out = out-of-frame oORF) **E.** The *IL2RA* pegRNA library designed to deliver TF motifs individually or in tandem was assayed in CD3+ primary T cells. Expression scores, normalised to control variants, are plotted. Variants are grouped by TF motif and error bars represent standard error across *n* = 2 replicates. (** = *q* < 0.01 and * = *q* < 0.05, Welsch’s unequal variances two-sample t-test against neutral variants with BH correction. **F.** Correlation (Spearman’s ρ) of expression scores tested in activated Jurkat and primary T cells for *n* = 77 variants from the library in (F), coloured by insertion type.

**Supplementary Figure 5.**
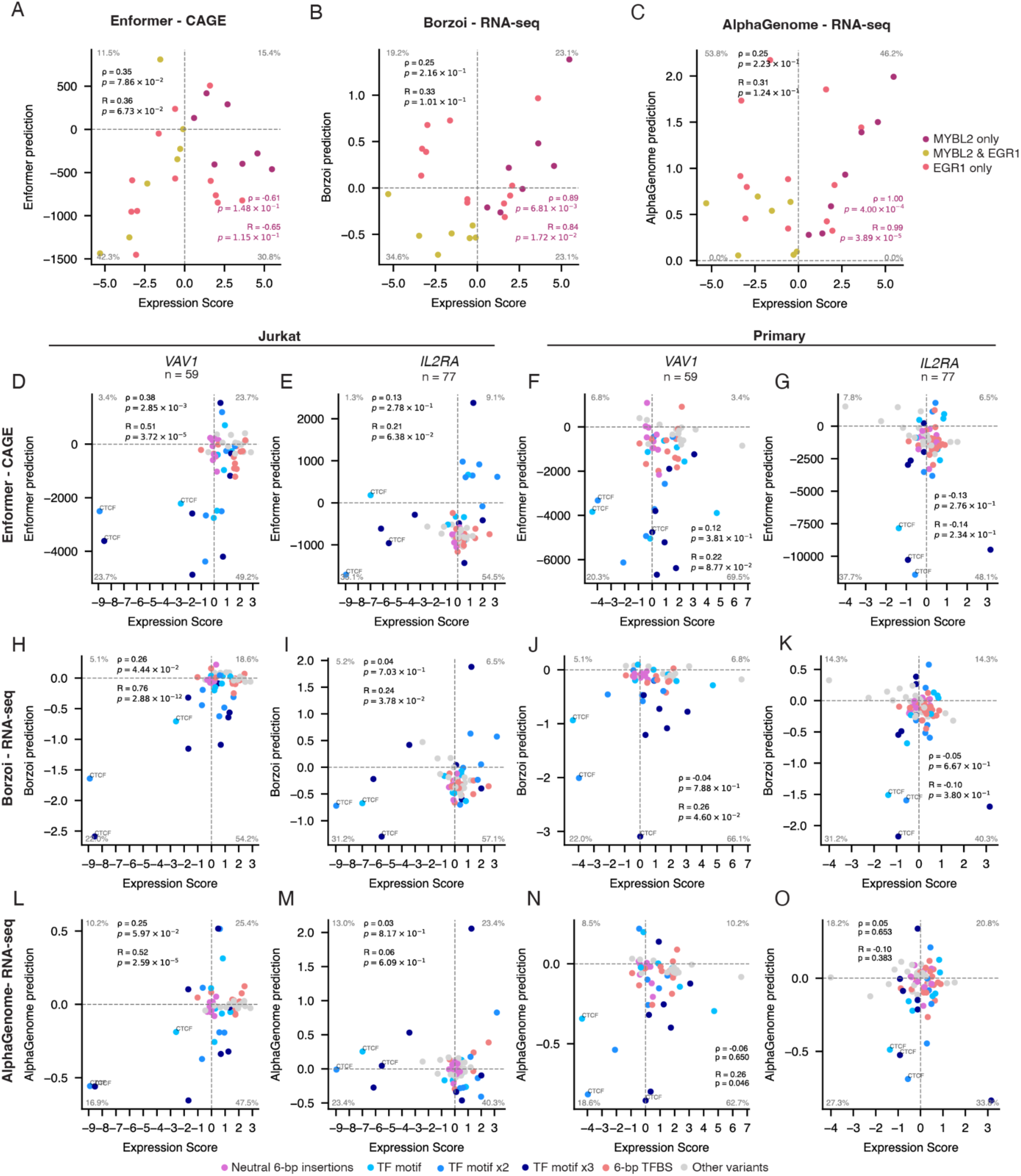
Exploring performance of sequence-to-expression models with PETRA. A-C. Correlations (Spearman’s ρ and Pearson’s R) between expression scores and cell type-specific Enformer, Borzoi, and AlphaGenome predictions are plotted for combinations of MYBL2 and EGR1 motifs inserted to the *IL2RA* 5’ UTR in Jurkats (*n* = 26 alleles). Alleles are coloured by the motifs present. Correlations for all alleles are in black, whereas those for alleles with only MYBL2 motifs are in violet. **D-O.** Correlations (Spearman’s ρ and Pearson’s R) between expression scores and cell type-specific Enformer, Borzoi, and AlphaGenome predictions are plotted for variants in TF tandem insertion libraries tested in Jurkat and primary T cells (*n* = 77 *IL2RA* variants; *n* = 59 *VAV1* variants). Variants are coloured by insertion type and CTCF motif insertions are labelled.

## SUPPLEMENTARY TABLES

**Supplementary Table 1.** Expression scores for 13,935 6-bp insertions scored with PETRA in Jurkat cells.

**Supplementary Table 2.** Expression scores for 6-bp insertions scored in validation libraries.

**Supplementary Table 3.** Analysis of uORFs from the 6-bp insertion screens.

**Supplementary Table 4.** TF motif scores derived from expression scores of 6-bp insertions in Jurkat cells.

**Supplementary Table 5.** Expression scores for *IL2RA* alleles edited with MYBL2 and EGR1 motif insertions in Jurkat cells.

**Supplementary Table 6.** Expression scores for *VAV1* alleles edited with the TF motif insertion library in Jurkat cells.

**Supplementary Table 7.** Expression scores for *IL2RA* alleles edited with the TF motif insertion library in resting and activated Jurkat cells.

**Supplementary Table 8.** Anti-CD25 flow cytometry analysis.

**Supplementary Table 9.** Expression scores for 6-bp insertions in *IL2RA* assayed in CD3+ primary T cells.

**Supplementary Table 10.** TF motif scores derived from expression scores for *IL2RA* 6-bp insertions in CD3+ primary T cells.

**Supplementary Table 11.** Expression scores for *IL2RA* alleles edited with the TF motif insertion library in CD3+ primary T cells.

**Supplementary Table 12.** Expression scores for *VAV1* alleles edited with the TF motif insertion library in CD3+ primary T cells.

**Supplementary Table 13.** Expression scores and AlphaGenome, Enformer and Borzoi predictions for 13,935 6-bp insertions scored with PETRA in Jurkat cells.

**Supplementary Table 14.** Expression scores and AlphaGenome, Enformer and Borzoi predictions for 6-bp insertions in *IL2RA* assayed in CD3+ primary T cells.

**Supplementary Table 15.** Expression scores and AlphaGenome, Enformer and Borzoi predictions for *IL2RA* alleles edited with MYBL2 and EGR1 motif insertions in Jurkat cells.

**Supplementary Table 16.** Expression scores and AlphaGenome, Enformer and Borzoi predictions for *VAV1* alleles edited with the TF motif insertion library in Jurkat and CD3+ primary T cells.

**Supplementary Table 17.** Expression scores and AlphaGenome, Enformer and Borzoi predictions for *IL2RA* alleles edited with the TF motif insertion library in Jurkat and CD3+ primary T cells.

**Supplementary Table 18.** Oligonucleotides used in this study.

**Supplementary Table 19.** pegRNA designs used in this study.

## Notes

### Competing Interest Statement

The authors have declared no competing interest.

### Summary of Updates

Figure 5 added including new comparisons of PETRA scores to predictions from computational models. Supplemental files updates.

## REFERENCES

1. Frangoul, H. et al. CRISPR-Cas9 gene editing for sickle cell disease and β-thalassemia. N. Engl. J. Med. 384, 252–260 (2021).

2. Lee, D. et al. A method to predict the impact of regulatory variants from DNA sequence. Nat. Genet. 47, 955–961 (2015).

3. Jaganathan, K. et al. Predicting Splicing from Primary Sequence with Deep Learning. Cell 176, 535–548.e24 (2019).

4. Zeng, T. & Li, Y. I. Predicting RNA splicing from DNA sequence using Pangolin. Genome Biol. 23, 103 (2022).

5. Avsec, Ž., et al. Effective gene expression prediction from sequence by integrating long-range interactions. Nat. Methods 18, 1196–1203 (2021).

6. Sasse, A. et al. Benchmarking of deep neural networks for predicting personal gene expression from DNA sequence highlights shortcomings. Nat. Genet. 55, 2060–2064 (2023).

7. Sasse, A., Chikina, M. & Mostafavi, S. Unlocking gene regulation with sequence-to-function models. Nat. Methods 21, 1374–1377 (2024).

8. Smith, C. & Kitzman, J. O. Benchmarking splice variant prediction algorithms using massively parallel splicing assays. Genome Biol. 24, 294 (2023).

9. ENCODE Project Consortium et al. Expanded encyclopaedias of DNA elements in the human and mouse genomes. Nature 583, 699–710 (2020).

10. Kerimov, N. et al. A compendium of uniformly processed human gene expression and splicing quantitative trait loci. Nat. Genet. 53, 1290–1299 (2021).

11. Abdellaoui, A., Yengo, L., Verweij, K. J. H. & Visscher, P. M. 15 years of GWAS discovery: Realizing the promise. Am. J. Hum. Genet. 110, 179–194 (2023).

12. Tewhey, R. et al. Direct Identification of Hundreds of Expression-Modulating Variants using a Multiplexed Reporter Assay. Cell 165, 1519–1529 (2016).

13. Kircher, M. et al. Saturation mutagenesis of twenty disease-associated regulatory elements at single base-pair resolution. Nat. Commun. 10, 3583 (2019).

14. Klein, J. C. et al. A systematic evaluation of the design and context dependencies of massively parallel reporter assays. Nat. Methods 17, 1083–1091 (2020).

15. Canver, M. C. et al. BCL11A enhancer dissection by Cas9-mediated in situ saturating mutagenesis. Nature 527, 192–197 (2015).

16. Fulco, C. P. et al. Systematic mapping of functional enhancer-promoter connections with CRISPR interference. Science 354, 769–773 (2016).

17. Korkmaz, G. et al. Functional genetic screens for enhancer elements in the human genome using CRISPR-Cas9. Nat. Biotechnol. 34, 192–198 (2016).

18. Gasperini, M. et al. A Genome-wide Framework for Mapping Gene Regulation via Cellular Genetic Screens. Cell 176, 377–390.e19 (2019).

19. Martin-Rufino, J. D. et al. Massively parallel base editing to map variant effects in human hematopoiesis. Cell 186, 2456–2474.e24 (2023).

20. Findlay, G. M. et al. Accurate classification of BRCA1 variants with saturation genome editing. Nature 562, 217–222 (2018).

21. Martyn, G. E. et al. Rewriting regulatory DNA to dissect and reprogram gene expression. Cell 188, 3349–3366.e23 (2025).

22. Tak, Y. E. et al. CRISPR PERSIST-On enables heritable and fine-tunable human gene activation. bioRxivorg 2024.04.26.590475 (2024) doi:10.1101/2024.04.26.590475.

23. Anzalone, A. V. et al. Search-and-replace genome editing without double-strand breaks or donor DNA. Nature 576, 149–157 (2019).

24. Schmidt, R. et al. CRISPR activation and interference screens decode stimulation responses in primary human T cells. Science 375, eabj4008 (2022).

25. Mathis, N. et al. Predicting prime editing efficiency and product purity by deep learning. Nat. Biotechnol. 41, 1151–1159 (2023).

26. Yan, J. et al. Improving prime editing with an endogenous small RNA-binding protein. Nature (2024) doi:10.1038/s41586-024-07259-6.

27. Chen, P. J. et al. Enhanced prime editing systems by manipulating cellular determinants of editing outcomes. Cell 184, 5635–5652.e29 (2021).

28. Blaeschke, F. et al. Modular pooled discovery of synthetic knockin sequences to program durable cell therapies. Cell 186, 4216–4234.e33 (2023).

29. Sample, P. J. et al. Human 5′ UTR design and variant effect prediction from a massively parallel translation assay. Nat. Biotechnol. 37, 803–809 (2018).

30. Wieder, N. et al. Differences in 5’untranslated regions highlight the importance of translational regulation of dosage sensitive genes. Genome Biol. 25, 111 (2024).

31. DepMap: The Cancer Dependency Map Project at Broad Institute. https://depmap.org/portal/depmap/.

32. Grant, C. E., Bailey, T. L. & Noble, W. S. FIMO: scanning for occurrences of a given motif. Bioinformatics 27, 1017–1018 (2011).

33. Rauluseviciute, I. et al. JASPAR 2024: 20th anniversary of the open-access database of transcription factor binding profiles. Nucleic Acids Res. 52, D174–D182 (2024).

34. Koeppel, J. et al. Prediction of prime editing insertion efficiencies using sequence features and DNA repair determinants. Nat. Biotechnol. 41, 1446–1456 (2023).

35. Barbadilla-Martínez, L., Klaassen, N., van Steensel, B. & de Ridder, J. Predicting gene expression from DNA sequence using deep learning models. Nat. Rev. Genet. 1–15 (2025).

36. Linder, J., Srivastava, D., Yuan, H., Agarwal, V. & Kelley, D. R. Predicting RNA-seq coverage from DNA sequence as a unifying model of gene regulation. Nat. Genet. 57, 949–961 (2025).

37. Avsec, Ž., et al. AlphaGenome: advancing regulatory variant effect prediction with a unified DNA sequence model. bioRxiv 2025.06.25.661532 (2025) doi:10.1101/2025.06.25.661532.

38. Zhu, F. et al. The interaction landscape between transcription factors and the nucleosome. Nature 562, 76–81 (2018).

39. Duttke, S. H. et al. Position-dependent function of human sequence-specific transcription factors. Nature 631, 891–898 (2024).

40. de Boer, C. G. & Taipale, J. Hold out the genome: a roadmap to solving the cis-regulatory code. Nature 625, 41–50 (2024).

41. Matharu, N. et al. CRISPR-mediated activation of a promoter or enhancer rescues obesity caused by haploinsufficiency. Science 363, eaau0629 (2019).

42. Doman, J. L., Sousa, A. A., Randolph, P. B., Chen, P. J. & Liu, D. R. Designing and executing prime editing experiments in mammalian cells. Nat. Protoc. 17, 2431–2468 (2022).

43. Nelson, J. W. et al. Engineered pegRNAs improve prime editing efficiency. Nat. Biotechnol. 40, 402–410 (2022).

44. Fornes, O. et al. JASPAR 2020: update of the open-access database of transcription factor binding profiles. Nucleic Acids Res. 48, D87–D92 (2020).

45. ENCODE Project Consortium. An integrated encyclopedia of DNA elements in the human genome. Nature 489, 57–74 (2012).

46. Mathis, N. et al. Machine learning prediction of prime editing efficiency across diverse chromatin contexts. Nat. Biotechnol. (2024) doi:10.1038/s41587-024-02268-2.

47. Whiffin, N. et al. Characterising the loss-of-function impact of 5’ untranslated region variants in 15,708 individuals. Nat. Commun. 11, 2523 (2020).

48. Arafeh, R., Shibue, T., Dempster, J. M., Hahn, W. C. & Vazquez, F. The present and future of the Cancer Dependency Map. Nat. Rev. Cancer 25, 59–73 (2025).

